# Expansion of *Drosophila* haemocytes using a conditional *GeneSwitch* driver affects larval haemocyte function, but does not modulate adult lifespan or survival from infection

**DOI:** 10.1101/2024.09.17.613448

**Authors:** Dan J Hayman, Lola M Morrin, Sudipta Halder, Eleanor J Phillips, Mirre J P Simons, Iwan R Evans

**Author notes:** Senior/joint corresponding authors.

## Abstract

Macrophages are responsible for diverse and fundamental functions in vertebrates. Fruit flies harbour an innate immune system of which the most populous blood cell (haemocyte) type bears striking homology to the vertebrate macrophage. The importance of these cells has been demonstrated previously, where immune and developmental phenotypes have been observed upon haemocyte ablation using pro-apoptotic transgenes driven by the *Hml* promoter.

Here we show that, as well as ablating *Hml*-positive cells *in vivo* using the pro-apoptotic transgene *bax*, we can also increase *Hml*-positive cell numbers using a constitutively-active form of *ras*. However, in adults, compared to larvae, total blood cell numbers were not significantly affected by experimental expansion or ablation. This therefore implies the existence of feedback mechanisms which regulate the number of haemocytes.

No effect on lifespan was observed from driving *ras* and *bax* in *Hml*-positive cells using a conditional genetic system (*Hml*-*GeneSwitch*). Using a constitutive driver system, we did observe differences in lifespan, however we attribute this to differences in genetic background that could have led to spurious conclusions. Additionally, no effect of either transgene was observed upon infection with two different bacterial species, although a striking pupal lethality phenotype was observed upon expansion of *Hml*-positive cells in the context of a self-encapsulation mutant genetic background. The latter confirms that the change in *Hml*-positive cell number does result in a phenotype. The lack of adult phenotypes could be due to the strength of our experimental manipulation or due to compensation via feedback mechanisms that operate to maintain total blood cell numbers. Our study demonstrates the importance of a conditional approach to modulate haemocyte cell numbers *in vivo* which allows for more precise study of innate immune system function. This approach could be especially fruitful to uncover the mechanisms that regulate total blood cell numbers across development and ageing.

## Introduction

Vertebrate macrophages enact diverse functions, including clearance of infections, removal of apoptotic corpses and tissue remodelling as part of development (Wynn et al. 2013). As a result of this diversity in function, macrophages are by necessity highly heterogeneous populations of cells, showing high levels of variation at both functional and transcriptomic levels (Murray et al. 2014; Colin et al. 2014; Gordon & Taylor 2005; Gordon et al. 2014). As a result of this heterogeneity, the study of macrophages in vertebrate systems is challenging, especially considering the necessity to also consider contributions from the adaptive immune system. Therefore the use of invertebrate models which represent vertebrate macrophages well is of great value to the field of immunology.

The fruit fly *Drosophila melanogaster* is such a model system. Flies lack an adaptive immune system and solely harbour an innate immune system, which consists of circulatory blood cells known as haemocytes (Buchon et al. 2014; Holz et al. 2003). Broadly, haemocytes consist of three distinct cell types, crystal cells, lamellocytes and plasmatocytes, the latter of which dominates the haemocyte population by quantity and bears considerable homology both functionally and transcriptomically to vertebrate macrophages (Yoon et al. 2023; Buchon et al. 2014; Wood & Jacinto 2007; Evans et al. 2003). These cells are able to migrate to wound sites, engulf pathogens and deposit extracellular matrix, all of which are essential characteristics of macrophages, as well as displaying transcriptomic and functional heterogeneity (Coates et al. 2021; Ratheesh et al. 2015). Considering these features of this class of *Drosophila* haemocyte, the fruit fly has been utilised frequently as a model of macrophages and associated behaviours (Ratheesh et al. 2015; Wood & Jacinto 2007).

Considering the wide array of roles *Drosophila* haemocytes are responsible for, it would seem logical for flies genetically engineered to be void of haemocytes to be inviable. Interestingly however, the developmental lethality of flies in which haemocytes are ablated appears to vary depending on the specific genetic driver used. Different studies have used different parts of the promoter region of the haemocyte-specific haemolectin (*Hml*) gene to genetically ablate haemocytes. Flies in which haemocytes are ablated using the *HmlΔ* driver complete development, albeit with an approximately halved ratio of successful eclosion from pupae relative to control flies, but this phenotype is rescued by germ-free or antibiotic-supplemented conditions (Shia et al. 2009; Charroux & Royet 2009; Arefin et al. 2015; Defaye et al. 2009; Ayyaz et al. 2015). In some studies melanotic spots were observed in flies in which haemocytes had been ablated, proposed to be the result of a lamellocyte expansion in *HmlΔ*-positive cell-ablated conditions (Defaye et al. 2009; Arefin et al. 2015). Comparatively, flies in which haemocytes are ablated using a different region of the *Hml* promoter, named *Hml*^P2A^, are severely pupal lethal, and are unable to be rescued (Stephenson et al. 2022). However, haemocyte ablation has not been investigated intensively with condition-dependent drivers. The benefit of conditional drivers is that all genetics are controlled for, with a genetic system conditionally activated by specific drugs (Clausen et al. 1999; Bockamp et al. 2008; Osterwalder et al. 2001) or environmental variables, such as temperature (McGuire et al. 2003; del Valle Rodríguez et al. 2011) and light (de Mena & Rincon-Limas 2020). A small number of experiments have used a temperature-sensitive Gal80 to ablate haemocytes conditionally (Defaye et al. 2009; Monticelli et al. 2024; Ayyaz et al. 2015), one of which revealed a reduced post-infection survival phenotype upon conditional haemocyte ablation (Defaye et al. 2009). Another showed that ablation of haemocytes impairs intestinal regeneration following oxidative stress-mediated damage or infection with the pathogens *Erwinia carotovora carotovora* strain 15 or *Pseudomonas entomophila* (Ayyaz et al. 2015). However, never before has a non-temperature-dependent conditional approach been used to ablate haemocytes. In addition, it has not been established whether haemocyte function is altered by increasing the number of cells; in other words, the literature suggests that a lack of haemocytes negatively affects haemocyte-driven functions, but does an excess of haemocytes produce the inverse of this?

Here we show the importance of the use of conditional drivers, especially for lifespan as a phenotype. Lifespan is highly polygenic making it critical to homogenise genetic backgrounds or use conditional systems where possible (Ford & Tower 2005). Our results differed between experiments using constitutive and conditional drivers, despite both systems using the same promoter region, *HmlΔ*. The constitutive *HmlΔ* driver (*HmlΔ*-*Gal4*) successfully expanded and ablated *Drosophila Hml-*positive haemocytes through UAS-regulated *ras85D^V12^* and *bax* transgenes, respectively, and this was correlated to an apparent modulation of lifespan. However, with our newly generated conditional driver, *HmlΔ*-*GeneSwitch*, no lifespan phenotype was observed, in spite of a modulation of *Hml*-positive cell numbers comparable to the constitutive driver experiments. Surprisingly, altering *Hml*-positive cell numbers conditionally did not change post-infection survival, raising the possibility that somehow the conditional modulation of *Hml*-positive haemocyte numbers did not change overall function. However, expansion of the *Hml*-positive cell population did modulate haemocyte function in larvae, which we demonstrated using a genetic model of self-encapsulation (Mortimer et al. 2021).

Bleeding larvae to obtain haemocyte populations revealed a large increase in overall haemocyte cell numbers when *HmlΔ*-*GeneSwitch* was used to drive *ras85D^V12^*.

However, in adult bleeds this was not the case, even though the *Hml*-positive population did grow in number *in vivo*. Conversely, our ablation conditions in adults did result in low numbers of *Hml*-positive cell numbers, but no change in overall bled cell numbers was observed. This suggests mechanisms that regulate total blood cell numbers could compensate for the ablation of *Hml*-positive haemocytes, despite strong initial manipulations in the larva. Our findings show that haemocyte numbers in larvae have functional relevance; more careful study of these functions *in vivo* will be possible using well-controlled conditional genetic systems such as presented herein.

## Methods

### Fly culture conditions

All experimental flies were kept at 25°C. Flies were mated for 2 days following eclosion, and were subsequently sorted into sexes and the desired genotypes for the experiment. Fly food consisted of the following components, as previously described (McCracken, Buckle, et al. 2020; McCracken, Adams, et al. 2020) : 8% yeast, 13% table sugar, 6% cornmeal, 1% agar and 0.225% nipagin (all w/v). In the case of growing bottles, 0.4% (v/v) propanoic acid was also added. Where the GeneSwitch construct was utilised, food was supplemented with 200 μM RU486 (final media concentration; dissolved in 8.6 mL absolute ethanol per 1 L of fly media and mixed into the media) or control food lacking RU486 (but still containing 8.6 mL absolute ethanol per 1 L of fly media). The food used for control and RU486 food was split from a single media preparation, meaning the exact same food was used for both conditions at the same time.

### *Drosophila* genotypes utilised

All fly genotypes utilised in this study are listed in Table 1, including where the line was obtained from, where applicable.

**Table 1.**
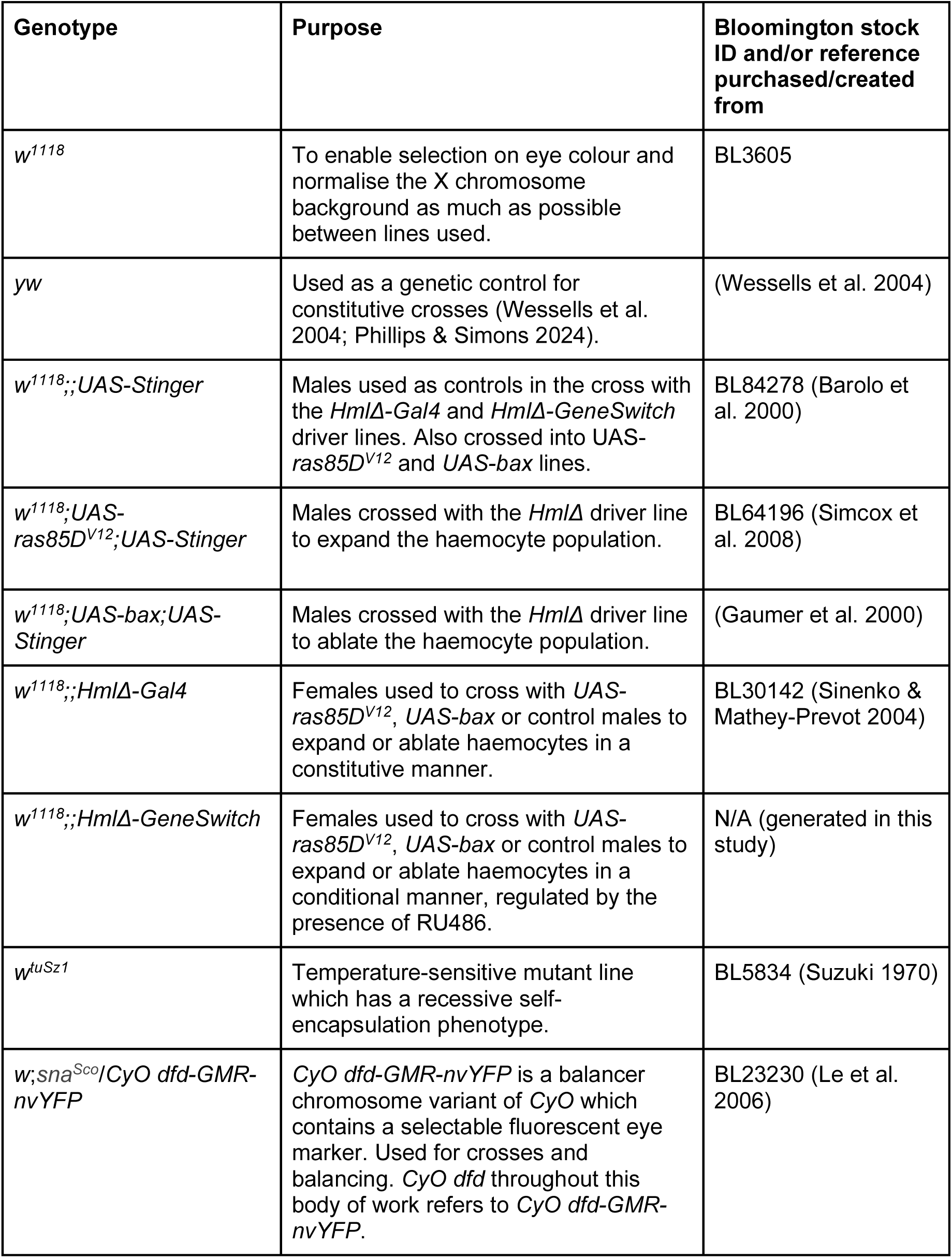
*Drosophila melanogaster* lines used in this research, specifying their use and source.

### Generation of *HmlΔ*-*GeneSwitch* flies

The *ELAV-GeneSwitch* plasmid (Osterwalder et al. 2001), a gift from Haig Keshishian (Addgene plasmid #83957; http://n2t.net/addgene:83957; RRID:Addgene_83957), was digested at 37°C for 1 hour with *Bgl*II and *Psp*XI (New England BioLabs), following the manufacturer’s standard protocol. Genomic DNA was isolated from 20 *Drosophila melanogaster w^1118^* strain flies, utilising the DNeasy Blood & Tissue Kit (QIAGEN), following the manufacturer’s standard protocol. The *HmlΔ* promoter was subsequently amplified from 80 ng of genomic DNA using Platinum SuperFi II Polymerase (Invitrogen), with the following primers (*Bgl*II and *Psp*XI restriction sites shown in lower case; additional bases added to maintain the GeneSwitch frame underlined), as previously used in the literature (Makhijani et al. 2011): F primer 5’-CGGTCACTcctcgaggCAAAAGTTATTTCTGTAGGC-3’; R primer 5’-CGGTAACTagatct<u>GGCTGCTGGGAGTCCAT</u>TTTGTTAGGCTAATCGGAAATTG-3’. The amplified *HmlΔ* promoter and digested *ELAV-GeneSwitch* plasmid were each run on a 1% (w/v) agarose TAE gel at 80 V for 1 hour and gel purified using the QIAquick Gel Extraction Kit (QIAGEN). Ligation of the insert and backbone was undertaken at 4°C for 16 hours followed by room temperature (22°C) for 2 hours, at a molar ratio of 10:1 (insert:backbone, using 50 ng insert DNA), performed using 1 U T4 DNA Ligase (Invitrogen), in a 20 µL reaction. TOP10 Competent Cells (ThermoFisher Scientific) were transformed with the ligation mix using the heat-shock method, recovered in a 37°C shaking incubator at 225 rpm for 1 hour, and plated on agar plates supplemented with carbenicillin (100 µg/mL), which were incubated at 37°C overnight. Discrete colonies were picked and following growth overnight in carbenicillin-supplemented LB broth at 37°C and 225 rpm, plasmid DNA was isolated using the QIAquick Spin Miniprep Kit (QIAGEN). The entire plasmid was sequenced using Oxford Nanopore long reads (Plasmidsaurus) and transformed into *w^1118^ Drosophila melanogaster* flies using P-element transposition (GenetiVision). Standard fly husbandry methodology was used to generate homozygous flies containing the *HmlΔ*-*GeneSwitch* construct, which were subsequently tested for their ability to drive the *UAS-Stinger* transgene in a haemocyte-specific manner.

### Imaging and quantification of haemocytes *in vivo*

Prior to imaging, adult flies were immobilised by incubation at -20°C for 10 minutes. An MZ205 FA fluorescent dissection microscope with a 1x PLANAPO objective lens (Leica) was subsequently used to image individual flies on the GFP channel (ED-GFP filter), using LasX image acquisition software (Leica). The following settings were used: exposure = 800 ms, gain = 1.0, zoom = 30X. Haemocyte numbers were quantified using the Find Maxima tool in Fiji (Schindelin et al. 2012), with Prominence set to 20. Differences between conditions were assessed using a two-way ANOVA. Tukey HSD post-hoc tests were used to specifically assess the statistical significance of differences relative to control flies.

To image haemocytes in larvae, wandering L3 larvae of the appropriate genotypes were selected and washed in distilled water and blotted dry. They were then imaged in distilled water on ice using the same microscope as above (2x PLANAPO objective lens, ED-GFP filter, exposure = 300 ms, zoom = 1.34, gain = 3.0).

### Assessment of post-infection survival

*Pseudomonas entomophila* (gift from Julia Cordero, University of Glasgow, UK) and *Staphylococcus aureus* (gift from Pedro Vale, University of Edinburgh, UK) bacteria (Vodovar et al. 2005; Perochon et al. 2021) were individually grown overnight at 29°C in LB broth, from stocks frozen at -80°C, to an OD600nm of 0.5. Flies were individually immobilised using a CO2 pad and a 0.15 mm diameter stainless steel insect minutien pin was washed in the bacterial stock, before being intrathoracically inserted into the immobilised fly for 2 seconds. Each fly was subsequently housed alone in a vial containing standard fly media (containing 8% (w/v) yeast) food, labelled with the time of infection and incubated in a temperature and humidity controlled room. The following day, flies were checked for death every 30 minutes from 1 hour after the light turned on. Cox proportional hazard models were used to test the effect of RU in the context of each genotype. The ‘coxph’ package in R was used to assess statistical significance (Therneau 2024).

### Longevity experiments

Flies were transferred to a custom cage to minimise disturbances, as previously described (McCracken, Adams, et al. 2020; Good & Tatar 2001). No more than 100 flies were housed in each cage. Following cageing, the number of deaths were recorded every other day, at which point dead flies were removed and fresh food provided, until no flies remained. In order to conditionally drive the *HmlΔ-GeneSwitch* construct throughout development, RU486-containing or control food was utilised from early larval stages. Proportional Cox hazard models were used between conditions in each genotype and each sex where applicable, including the ’cage’ as a random effect to correct for pseudoreplication within cages (McCracken, Buckle, et al. 2020; Therneau et al. 2003; Ripatti & Palmgren 2000). The ‘coxme’ package in R was used to assess statistical significance (Therneau 2012).

### Assessment of survival and melanisation in the context of the *tuSz^1^* mutation

Virgin females containing the *tuSz^1^* mutation (*w^tuSz1^*) were crossed with *w^1118^;If/UAS- ras85D^V12^;HmlΔ-GeneSwitch* males at 25°C on food containing either RU486 or control food. After 24 hours at 25°C, parents were removed and the vial containing eggs switched to either 29°C to enable the temperature-sensitive *tuSz^1^* melanotic phenotype to manifest, or kept at 25°C as negative controls. To assess if expansion of haemocytes caused lethality, offspring on both diets at both temperatures were sexed and counted, as well as being genotyped using the visible marker *If* to discriminate the presence of *UAS-ras85D^V12^*. At the L3 stage of development, 27 male larvae on a control diet and 25 male larvae on an RU-containing diet at 29°C were randomly picked and their melanisation level scored from 1-5, based on a novel, semi-quantitative scoring system (shown in Supplementary Figure 6). Each larva was individually housed in a fresh vial and eventually checked for genotype upon eclosion. Flies which did not eclose were checked for their genotype, via *If*, by removing the puparium. Larvae were imaged on a MZ205 FA fluorescent dissection microscope with a 1x PLANAPO objective lens (Leica) on the brightfield channel, using LasX image acquisition software (Leica), with the following settings: exposure = 700 ms, gain = 1.0 and zoom = 40X. To image in detail the fat body of developmentally arrested flies following puparium removal, the same microscope was utilised, but with zoom set to 120X.

### Dissection, staining, imaging and analysis of larval and adult *ex vivo* bleeds

Wandering L3 larvae of the appropriate genotype were selected and washed in distilled water, blotted dry and then transferred to 75 *μ*m of S2 media [Schneiders medium (Merck) supplemented with 1X Pen/Strep (Gibco) and 10% heat-inactivated FBS (Gibco)] on ice in a petri dish. Larvae were opened using size 5 forceps from their posterior spiracles towards the anterior end to release haemocytes into S2 media and then agitated for 10 seconds to release more tightly-adhered cells. The 75 *μ*l of S2 media containing haemocytes was then transferred to a 96-well plate (Greiner) and a further 75 *μ*l of S2 media added. For adult bleeds, flies were placed at -20°C for 3 mins ahead of dissection to stop movement. Two newly-eclosed adult flies (day one) of the same sex were then dissected in a single droplet of 75 *μ*l droplet of S2 media on ice. Flies were decapitated and then cut longitudinally along their head, thorax and abdomen within that droplet and then an additional 75 *μ*l of S2 media added. Carcasses were agitated ten times via pipetting before cell suspensions were transferred to a well in a 96-well plate. For both larval and adult bleeds, cells were allowed to adhere at room temperature and in the dark for 45-60 mins before fixation.

Cells adhered to wells were fixed in 4% formaldehyde (diluted from 16% EM grade, Thermofisher) solution in PBS for 15 mins, before permeabilisation using 0.1% Triton-X100 (Sigma) in PBS (Oxoid) for 5 mins. Cells were stained using either TRITC-phalloidin (1:1000, Invitrogen; all dissections with the exception of *HmlΔ- Gal4*, constitutive L3 larvae) or Texas Red-Phalloidin (1:400, Invitrogen; *HmlΔ-Gal4*, constitutive L3 larvae) along with nucBlue nuclear stain (2 drops per mL, Invitrogen).

PBS washes were performed after each step and cells were imaged in a final volume of 200 *μ*L PBS using an MDX ImageExpress Hi-Content microscope (10X lens).

Dapi, Cy3, Texas Red and GFP filter sets were used to image nucBlue, TRITC-phalloidin, Texas Red-phalloidin and Stinger fluorescence, respectively. Nine sites were imaged per well.

Images from at least 6 fields of view (FOV) per well were exported as Tiff files using ImageExpress software. For larval bleeds, cell counts per FOV were made from the Dapi/nucBlue channel images. These images were segmented (thresholded, fill holes, watershed) to create binary images, which were then counted automatically using the analyse particles tool in Fiji. All contrast adjustment and thresholding were applied equally to each Tiff file within each coherent data set.

For adult dissections, contrast adjustments were applied to Tiff images for each channel to create an RGB merged image; as previously each channel was treated equally across experimental groups. Using the point selection tool and channels tool in Fiji the number of *Hml*-positive cells with a nucBlue-stained nuclei were counted in the green/EGFP and Dapi channels; *HmlΔ-Gal4* or *HmlΔ-GeneSwitch* were used to drive expression from *UAS-Stinger* in these samples. Non *Hml*-positive haemocytes were then scored on the basis of cell morphology (F-actin staining via phalloidin/red channel fluorescence) and the presence of nucBlue staining. Cells without nucBlue staining were excluded.

Statistical significance was calculated using ANOVAs for the conditional driver, with pairwise comparisons assessed using Tukey’s HSD post-hoc testing, or Student’s two-tailed t-tests for constitutive data.

## Results

### Constitutive and conditional *HmlΔ* drivers are able to expand or ablate *Hml*-labelled haemocyte populations

We used *HmlΔ*-*Gal4* to constitutively drive either the pro-apoptotic gene *bax*, a potent activator of apoptotic pathways in the fly (Gaumer et al. 2000), in order to ablate *HmlΔ*-positive cells, or *ras85D^V12^*, a constitutively-active mutant variant of *ras* used to drive expansion of the *HmlΔ*-positive cell population (Hobbs et al. 2016; Ashburner et al. 1999; Simcox et al. 2008). Expression from *UAS-Stinger*, which encodes a nuclear-localised GFP (Barolo et al. 2000) enabled labelling of *Hml*- positive cells. Virtually no *Hml*-positive cells were visible in flies where *HmlΔ-Gal4* was used to drive *UAS-bax*, as found previously (Defaye et al. 2009), and increased numbers of *Hml*-positive cells were observed in flies where *HmlΔ*-*Gal4* was used to drive *UAS-ras85D^V12^* (Figures 1A and 1B). We observed a slight increase in the developmental lethality of flies carrying *HmlΔ-Gal4* and either *UAS-bax* or *UAS- ras85D^V12^*, relative to control flies only carrying *HmlΔ-Gal4* (Figure 1C). The increase in mortality in flies carrying *UAS*-*bax* flies achieved statistical significance (χ^2^1 statistic = 6.43, p-value = 0.011), whereas the apparent increase in *UAS-ras85D^V12^* flies was not significant (χ^2^1 statistic = 1.40, p-value = 0.236).

**Figure 1.**
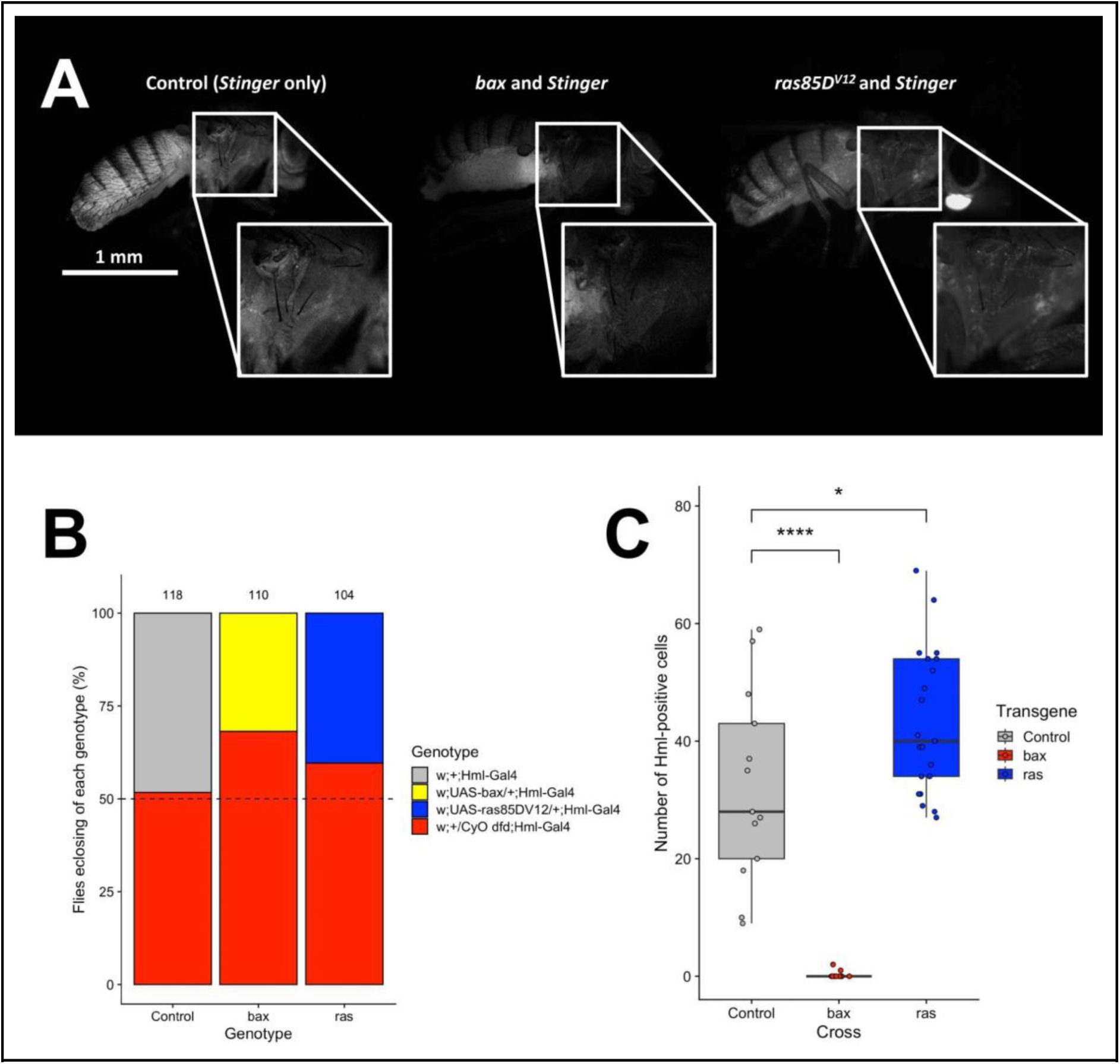
*HmlΔ*-*Gal4* can be used to expand or reduce numbers of *Hml*-labelled cells. (**A**) Driving *bax* or *ras85D^V12^* via *HmlΔ-Gal4* both produced viable progeny and modulated numbers of *HmlΔ*-labelled cells in female flies; *UAS*-*bax* reduced the labelled cell number to almost no cells, whereas *UAS-ras85D^V12^* increased the number of cells when imaged at 1-day post-eclosion. Cells labelled via expression from *UAS-Stinger*; Stinger is a nuclear-localised GFP, which enables visualisation of *HmlΔ*-positive cells. (**B**) Female *w^1118^;;HmlΔ-Gal4* flies were crossed with male flies of each of the following genotypes: *w^1118^;+/CyO dfd* (’Control’), *w^1118^;UAS-bax/CyO dfd* (’bax’) and *w^1118^;UAS-ras85D^V12^/CyO dfd* (’ras’). The number of flies eclosing with *CyO dfd* was compared to those eclosing without it, so as to genotype the eclosing flies. The estimated ratio for each genotype was 50%, indicated by the dotted line, and each genotype was compared to the ratios observed in ’Control’ flies using a χ^2^ test. The total number of progeny counted from each cross is shown above each bar. (**C**) Raw cell numbers detected via the Find Maxima tool from superficial, lateral images of adult flies at 1- day post-eclosion for each transgene are shown. A one-way ANOVA with Tukey HSD post-hoc tests was used to calculate statistical significance, for which * and **** signifies adjusted p-values ≤ 0.05 and 0.0001, respectively. The exact genotypes used in panels A and B were as follows: ’Control’ = *w^1118^;;UAS-Stinger/HmlΔ-Gal4*; ’ras’ = *w^1118^;+/UAS-ras85D^V12^;UAS-Stinger/HmlΔ-Gal4*; ’bax’ = *w^1118^;+/UAS-bax;UAS-Stinger/HmlΔ-Gal4*.

*HmlΔ-Gal4* is a constitutive driver and leads to the expression in differentiated blood cells from larval stages onwards. This approach however requires the crossing of different genetic lines that are unlikely to be genetically identical. Conditional drivers have the benefit of controlling fully for genotype, therefore we generated a *HmlΔ*- *GeneSwitch* transgenic line. The *HmlΔ*-*GeneSwitch* construct we made resulted in strong expression of a *UAS-Stinger* transgene by 1-day post-eclosion, when activated by RU486 (RU) throughout development (Supplementary Figure 1). Under both ablated and expanded conditions (*HmlΔ-GeneSwitch* driving *UAS-bax* and *UAS-ras85D^V12^*, respectively, with RU provided throughout development and continuing post-eclosion), a modulation of cell number was detected with both transgenes at the tested time point of 6-days post-eclosion (Figure 2), demonstrating *HmlΔ*-*GeneSwitch* can fully reproduce results obtained via *HmlΔ-Gal4* (Figure 1).

**Figure 2.**
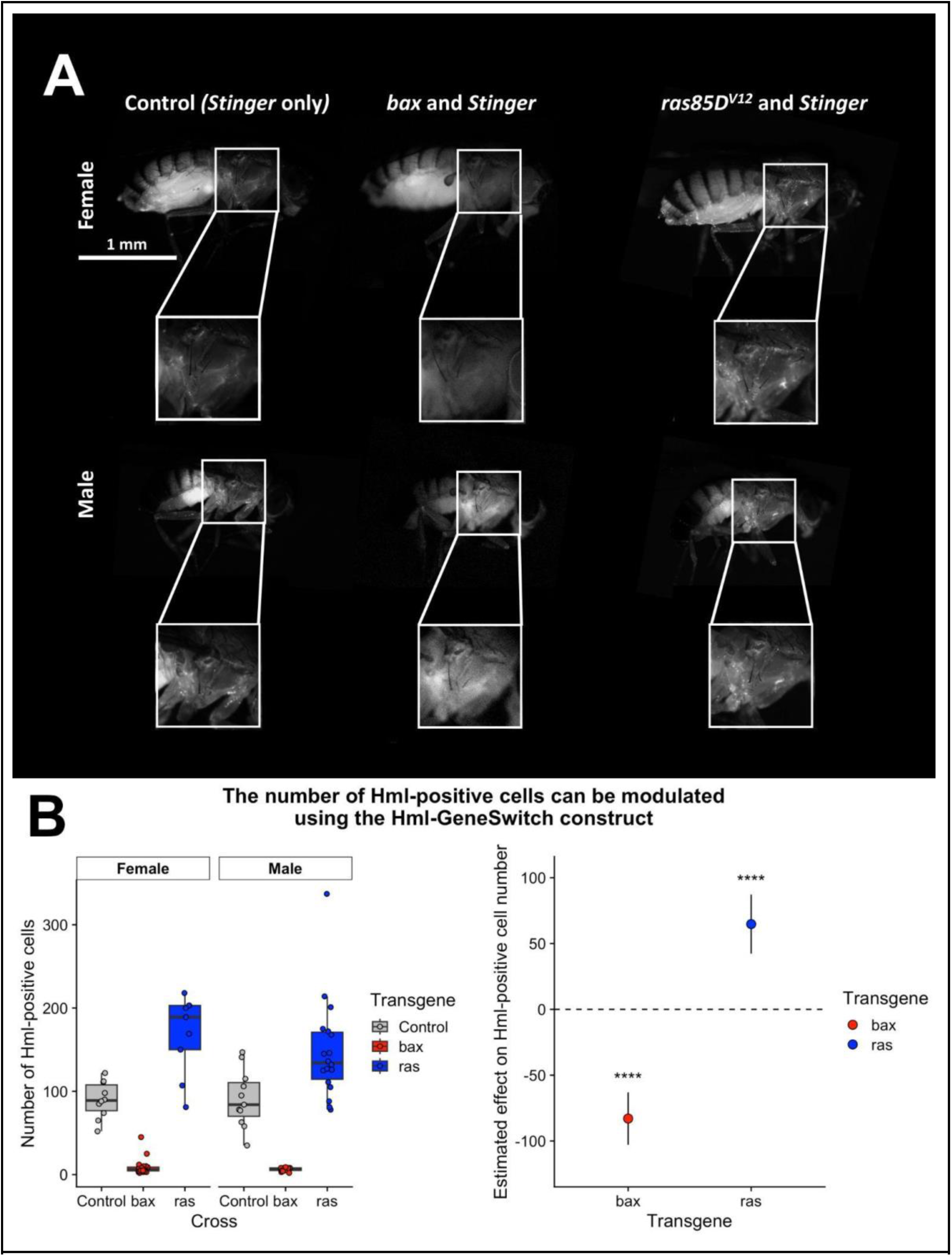
The *HmlΔ*-*GeneSwitch* construct can be induced by RU486 and is able to modulate numbers of *Hml*-labelled cells. (**A**) Driving *bax* or *ras85D^V12^* with *HmlΔ*- GeneSwitch both produced viable progeny when RU486 was supplied continuously from embryonic stages onwards. Representative images of flies at 6-days post-eclosion that are expressing *Stinger* only (‘Control’), *bax* and *Stinger*, or *ras85D^V12^* and *Stinger* (left to right). (**B**) Counting of *HmlΔ*-positive cells at 6-days post-eclosion using the Find Maxima tool on Fiji identified an increase in *HmlΔ*-positive cells in flies where *HmlΔ*-*GeneSwitch* was driving *ras85DV12* and a strong reduction of labelled cells when *bax* was driven. A two-way ANOVA with post-hoc Tukey HSD corrections was used to assess statistical significance relative to controls, as well as obtain 95% confidence intervals for the effect size of each transgene. **** indicates adjusted p-values < 0.0001. N ≥ 9 flies per condition. The exact genotypes used in this figure were: ’Control’ = *w1118*;+;*UAS-Stinger*/*HmlΔ-GeneSwitch*; ’bax’ = *w1118*;+/*UAS- bax*;*UAS-Stinger*/*HmlΔ-GeneSwitch*; ’ras’ = *w1118*;+/*UAS*-*ras85DV12*;*UAS-Stinger*/*HmlΔ- GeneSwitch*.

Note that our ability to visualise cells, using *UAS-Stinger*, requires induction of this system, thus we only visualise cells that have an active *HmlΔ* promoter.

Bleeding of larvae and adults allows for measuring total numbers of blood cells (including cells not labelled by *UAS-Stinger*), with the caveat that it does not capture cells tightly integrated into tissue or strongly adhered, which may therefore be harder to release during dissections. Despite this limitation, imaging of L3 larvae ahead of dissection demonstrated a strong enhancement of numbers of *Hml*-labelled cells compared to controls on constitutive expression of *ras85D^V12^* via *HmlΔ-Gal4*, whilst almost no cells were visible on expression of *bax* (Supplementary Figure 2A-C).

However, larval bleeds from the same genotypes revealed significant increases in total haemocyte numbers via both strategies (Supplementary Figure 2D-G). Very few Stinger-positive cells survived on expression of *bax* and many of those cells exhibited a lamellocyte-like morphology, whereas almost all cells present on *ras85D^V12^* expression were Stinger-positive (Supplementary Figure 2E-F).

Bleeding of adult flies revealed that a significant difference was only achieved for flies in which *bax* was driven using constitutive *HmlΔ-Gal4*, with only a mild reduction in haemocytes observed (and then only in females). Furthermore, driving *ras85D^V12^* did not significantly expand numbers of bled cells (Supplementary Figure 2H-I). As per larvae, many cells present in adult bleeds on expression of *bax* exhibited lamellocyte-like morphologies (data not shown).

Conducting the equivalent experiments using *HmlΔ*-*GeneSwitch* revealed very similar results to the constitutive approaches (Supplementary Figure 3): in *HmlΔ*- *GeneSwitch* larvae driving *ras85D^V12^*, the number of cells was increased by more than 30-fold, whilst larvae in which *bax* was driven showed a trend towards an increase in cell number, although this did not achieve statistical significance (p-value = 0.06; Supplementary Figure 3A-E).

In adults where transgenes were driven by *HmlΔ*-*GeneSwitch*, no statistically significant changes in total cell numbers were observed (Supplementary Figure 3F), although the number of *Hml*-positive cells (interpreted from GFP-labelling) was significantly increased in flies where *ras85D^V12^* was driven by *HmlΔ*-*GeneSwitch*, relative to those in which *bax* was driven (Supplementary Figure 3G).

These data suggest the existence of limitations or feedback mechanisms that control numbers of haemocytes that can be present within adults, given that, despite dramatic increases in cell numbers in larvae, the effect on numbers of haemocytes in adults is less pronounced on expression of *ras85D^V12^*. Potentially, rather than modulating the number of total haemocytes, our genetic tools may in fact specifically modulate numbers of *Hml*-positive cells.

### Manipulating haemocyte cell numbers does not affect adult lifespan

Considering that immune function is central to ageing (Stanley et al. 2023), we wanted to investigate whether manipulating the haemocyte population would impact longevity. Upon testing this with large sample sizes under highly controlled conditions, we observed alterations to lifespan upon both *Hml*-positive cell expansion and ablation using constitutive expression of *ras85D^V12^* or *bax* via *HmlΔ-Gal4* (Supplementary Figure 4). However, this effect was not replicated when repeating this using the conditional *HmlΔ-GeneSwitch* system and continuous application of RU, despite ensuring that all other aspects of the experiment were identical (Figure 3; statistics shown in Supplementary Table 1). This strongly suggests that small genetic differences between the *UAS*-lines explained the difference in longevity observed via the constitutive *HmlΔ-Gal4* approach.

**Figure 3.**
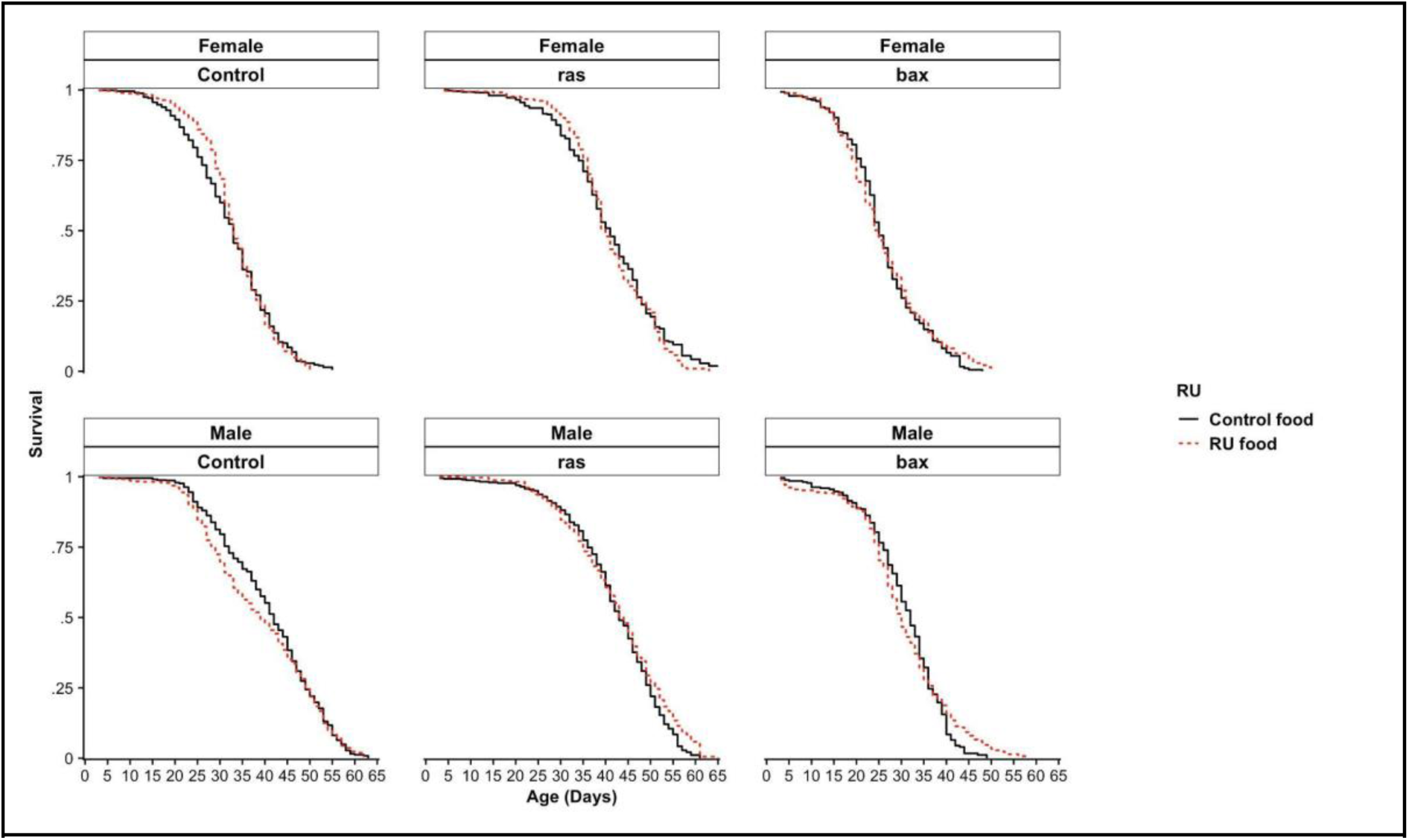
Neither ablation nor expansion of *HmlΔ*-positive cells significantly affects lifespan using the conditional *HmlΔ*-*GeneSwitch* driver. *HmlΔ*-*GeneSwitch* was activated from larval stages using food supplemented with RU486, and following eclosion flies were mated for 48 hours. Subsequently flies were sorted into lifespan cages, for which new food was provided every 48 hours and the number of dead flies was counted. For ’Control’ flies, *HmlΔ*-*GeneSwitch* drove only the *UAS*-*Stinger* transgene, whereas for ’bax’ and ’ras’ flies, either *UAS*-*bax* or *UAS*-*ras85D^V12^* were driven in addition to *UAS*-*Stinger*. The exact genotypes used above were as follows: ’Control’ = *w^1118^*;+;*UAS*-*Stinger*/*HmlΔ*-*GeneSwitch*; ’ras’ = *w^1118^*;+/*UAS*-*ras85D^V12^*;*UAS*-*Stinger*/*HmlΔ*-*GeneSwitch*; ’bax’ = *w^1118^*;+/*UAS*-*bax*;UAS- *Stinger*/*HmlΔ*-*GeneSwitch*.

Indeed, similarly *bax*, *ras85D^V12^* and control crosses with the conditional driver showed the same lifespan differences; the *bax* crosses lived shortest whilst the *ras85D^V12^* crosses lived longest, but with no additional modulation by driving the ablation or expansion using RU (Figure 3). Additionally, to test this hypothesis further, we crossed the same *bax*, *ras85D^V12^* and control lines to a *yw* background and found similar lifespan differences to those observed when using the constitutive *HmlΔ*-*Gal4* driver, particularly for the reduced lifespan observed with *bax* (Supplementary Figure 5). These observations fit with the idea that lifespan is a highly polygenic trait and that the accumulation of deleterious mutations through drift can strongly determine lifespan (Cui et al. 2019), underscoring the importance of conditional driver systems, especially when studying ageing (Ford & Tower 2005). It remains possible that stronger enhancement of haemocyte numbers could impact lifespan, while the non-*Hml* positive haemocytes that appear expanded upon expression of *bax* could mask effects on lifespan linked to compromised immunity.

We investigated whether *Hml*-positive cell number affects survival from a pathogen infection, as previously reported using constitutive drivers (Defaye et al. 2009; Charroux & Royet 2009), and including a small number of experiments using a temperature-sensitive conditional system (Defaye et al. 2009). Immune challenges with two commonly-used pathogens used to study immunity in the fly (*Pseudomonas entomophila* or *Staphylococcus aureus*) were not affected by conditional *Hml*- positive cell ablation or expansion (Figure 4; statistics shown in Supplementary Table 2). These results therefore imply that neither lifespan nor immune activity are severely impacted by modulating the number of *Hml*-positive cells. Again, more significant expansion of *Hml*-positive cells or the presence of increased numbers of *Hml*-negative haemocytes could obscure immune phenotypes in these assays.

**Figure 4.**
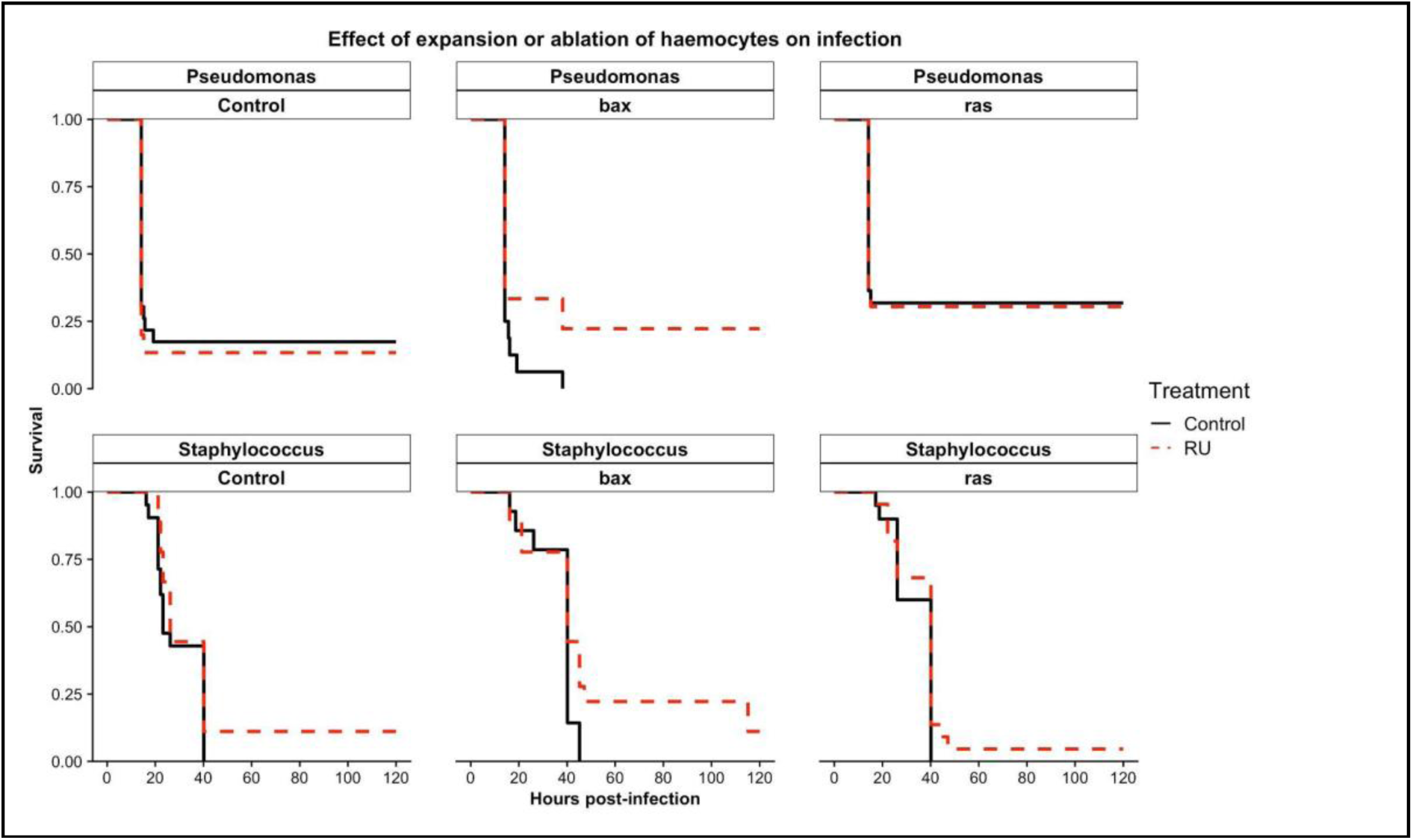
Conditional manipulation of *Hml*-positive number does not significantly affect post-infection survival with *Pseudomonas entomophila* or *Staphylococcus aureus* when using the conditional *HmlΔ*-*GeneSwitch*. Female flies containing *HmlΔ*-*GeneSwitch* and the transgenes *UAS*-*bax*, *UAS*-*ras85D^V12^* or *UAS*-*Stinger* (the control, all flies tested also possessed this transgene) were either activated or not in female flies by the presence of RU486 throughout development and infected with *Pseudomonas entomophila* or *Staphylococcus aureus* at 5-days post-eclosion. No statistically significant modulation of post-infection survival was observed upon infection with either bacterium (statistics shown in Supplementary Table 2). Infected flies were checked for deaths every 30 minutes from approximately 18-hours post-infection until approximately 25-hours post-infection, and less frequently after this point. Experiments were terminated after 120-hours post-infection. No flies ’infected’ with a sham infection (water) died over the 120-hour time course (not shown on plot). N ≥ 14 for each treatment group. The exact genotypes used above were as follows: ’Control’ = *w^1118^*;+;*UAS*-*Stinger*/*HmlΔ*-*GeneSwitch*; ’bax’ = *w^1118^*;+/*UAS*-*bax*;*UAS*-*Stinger*/*HmlΔ*- *GeneSwitch*; ’ras’ = *w^1118^*;+/*UAS*-*ras85D^V12^*;*UAS*-*Stinger*/*HmlΔ*-*GeneSwitch*.

### Modulation of *Hml*-positive cell number using *HmlΔ*- *GeneSwitch* affects the haemocyte-dependent phenotype of *tuSz^1^*

Using the *HmlΔ-GeneSwitch* driver we generated, we were able to change the number of *Hml*-labelled cells, but this was not associated with infection or lifespan phenotypes. Therefore, we wanted to test another known haemocyte-related function upon manipulation of their numbers, as a positive control. To do this, we used the *tuSz^1^* mutant; a mutation on the X chromosome, 35 bp upstream of the *GCS1* transcription start site. This mutation results in a loss of self-tolerance in the posterior fat body and a haemocyte-driven self-encapsulation phenotype that is temperature dependent (Mortimer et al. 2021). We hypothesised that expansion of the haemocyte cell population would lead to increased targeting of misrecognised ’self’ tissue in the presence of the *tuSz^1^* mutation (Mortimer et al. 2021). Indeed, we observed developmental lethality when *ras85D^V12^* was overexpressed using *HmlΔ*- *GeneSwitch*, with lethality stronger at 29°C (the temperature at which the melanisation phenotype presents itself in control flies). In males at the higher temperature, where the *tuSz^1^* mutation cannot be rescued by a second copy of the X chromosome, near-complete developmental lethality was observed (Figure 5A and Table 2); only a single fly eclosed at 29°C where *UAS*-*ras85D^V12^* had been activated by *HmlΔ*-*GeneSwitch* and this fly died within 3 days of eclosion. Flies that overexpressed *ras85D^V12^* alone at 29°C, without the *tuSz^1^* mutation, showed no developmental lethality (Figure 5B; and Table 2). Note, we attempted similar experiments with *HmlΔ*-mediated expression of *UAS-bax*, but these experiments failed due to genetic incompatibilities between stocks. Flies overexpressing *ras85D^V12^* in the same genetic background but at 25°C also showed no developmental lethality (Supplementary Figure 7 and Table 2), as well as the 25°C RU-driven flies showing no significant difference to flies grown at 25°C where *ras85D^V12^* was driven by the constitutive *HmlΔ*-*Gal4* driver (Table 2, counts from Figure 1C; these flies were female only so as to match the constitutive counts). This therefore suggested that there is no difference in effect on lethality of *UAS*-*ras85D^V12^* between the constitutive and conditional *HmlΔ* drivers.

**Figure 5.**
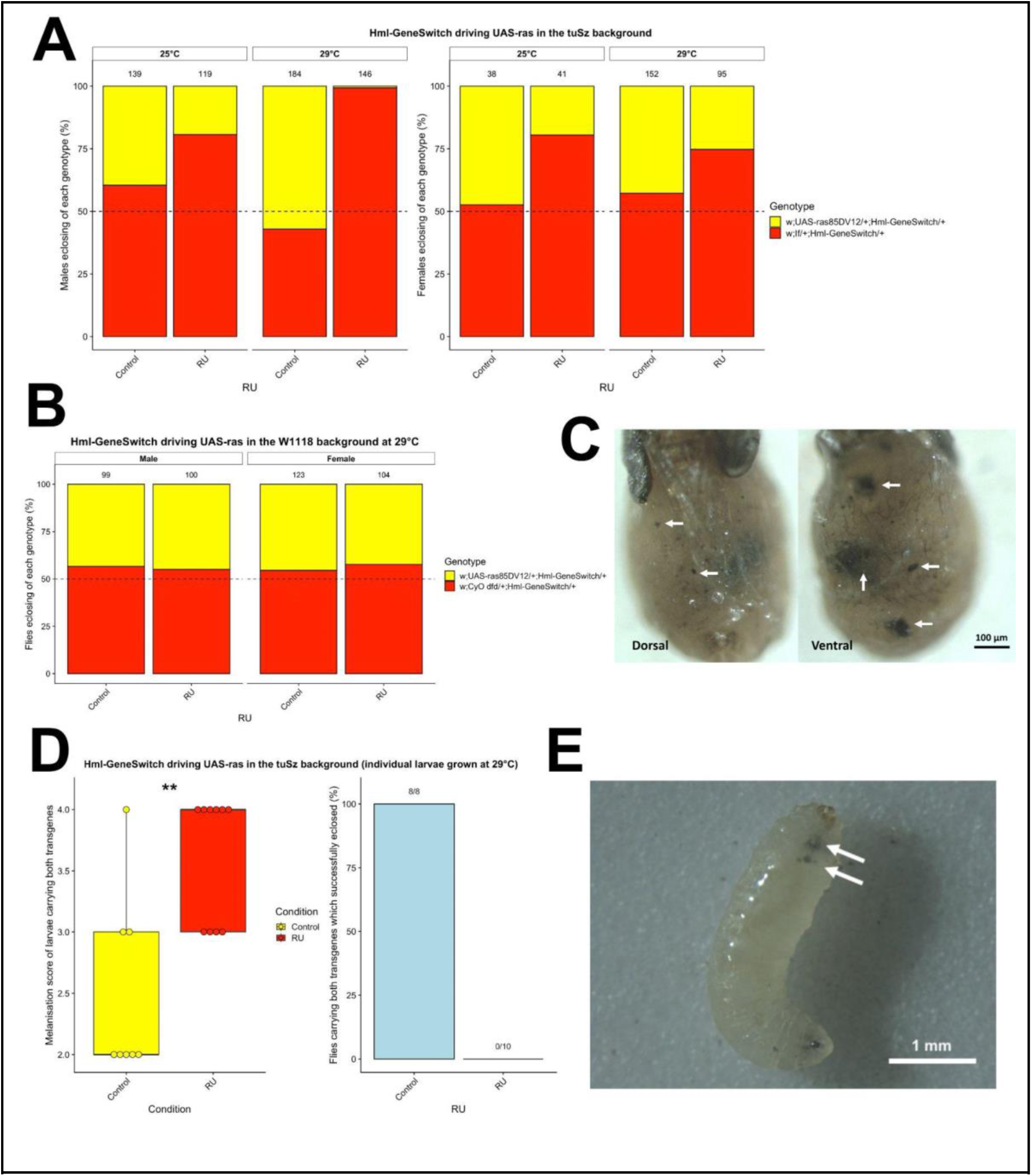
Driving *UAS*-*ras85D^V12^*with *HmlΔ*-*GeneSwitch* causes pupal lethality as a result of excessive melanotic tissue. (**A**) Far fewer male flies eclose with both the *UAS*- *ras85D^V12^* and *HmlΔ*-*GeneSwitch* transgenes when grown at 29°C in the presence of RU than female flies in any condition, or males in the absence of RU or grown at 25°C. The *If* marker was used to identify whether flies enclosed with both transgenes or with only *HmlΔ*- *GeneSwitch*, and therefore the expected ratio of each genotype was 50% (indicated by the dotted line). The numbers above each bar signify how many total eclosed flies were counted per condition. The cross to generate these flies was as follows: w*^tuSz1^* (female containing the *tuSz^1^* mutation) X *w^1118^*;*If*/*UAS*-*ras85D^V12^*;*HmlΔ*-*GeneSwitch* (male). (**B**) Driving *UAS*- *ras85D^V12^* with *HmlΔ*-*GeneSwitch* in the absence of the *tuSz^1^* mutation did not result in developmental lethality at 29°C in either males or females. The dotted line indicated the expected ratio of flies carrying both the *HmlΔ*-*GeneSwitch* and *UAS*-*ras85D^V12^* trangenes, which was 50%. The flies crossed here to test developmental lethality were of the following genotypes: *w^1118^*;;*HmlΔ*-*GeneSwitch* (female) X *w^1118^*;*UAS*-*ras85D^V12^*/*CyO dfd* (male). (**C**) Male flies of the genotype *wtuSz1*;*UAS*-*ras85DV12*;*HmlΔ*-*GeneSwitch* grown at 29°C in the presence of RU arrested at a late pupal stage. Upon removal of the puparium, significant levels of melanotic tissue in the fat body were observed. Large patches of this are signified with white arrows. (**D**) Male L3 larvae from the same experiment were scored for levels of melanisation using the novel scoring system (Supplementary Figure 6). These larvae were followed individually throughout development so as to confirm their genotype upon eclosion. Of the flies with both *UAS*-*ras85DV12* and *HmlΔ*-*GeneSwitch*, flies fed food containing RU showed a statistically significant increase in melanisation relative to those fed a control diet. A two-sided Student’s t-test was used to assess statistical significance, for which ** signifies p-value < 0.01. 27 male larvae on a control diet and 25 male larvae on an RU-containing diet were originally picked from each condition, of which 8 control diet-fed larvae and 10 RU-fed harboured both *UAS*-*ras85DV12* and *HmlΔ*-*GeneSwitch*. All 8 of the control-fed larvae successfully pupated and eclosed, whereas none of the 10 RU-fed larvae eclosed, and instead aborted development at approximately the P12 pupal stage, as previously observed. (**E**) Example larva of the genotype *wtuSz1*;*UAS*-*ras85DV12*/+;*HmlΔ*-*GeneSwitch*/+ with clear excess melanisation visible, highlighted by white arrows.

**Table 2.** *χ*^2^ statistics for each eclosion comparison. A 2x2 contingency table was used for each comparison.

| Comparison | Genetic background | $\chi^2_1$ statistic | P-value |
| --- | --- | --- | --- |
| Male 25°C RU vs control | *tuSz*<sup>1</sup> | 12.45 | 0.00042 |
| Female 25°C RU vs control | *tuSz*<sup>1</sup> | 6.93 | 0.0085 |
| Male 29°C RU vs control | *tuSz*<sup>1</sup> | 118.68 | < 0.00001 |
| Female 29°C RU vs control | *tuSz*<sup>1</sup> | 7.77 | 0.0053 |
| Male 29°C RU vs control | *w*<sup>1118</sup> | 0.049 | 0.82 |
| Female 29°C RU vs control | *w*<sup>1118</sup> | 0.24 | 0.63 |
| Male 25°C RU vs control | *w*<sup>1118</sup> | 0.001 | 0.97 |
| Female 25°C RU vs control | *w*<sup>1118</sup> | 0.0009 | 0.98 |
| Female 25°C *Hml*Δ-*GeneSwitch* RU vs *Hml*Δ-*Gal4* RU | *w*<sup>1118</sup> | 0.84 | 0.36 |

Even through the puparium, it was clear that the flies which failed to eclose showed high levels of excessive melanisation (Supplementary Figure 8). Removal of the puparium revealed that these pupal-lethal flies consistently died at approximately the P12 pupal stage (Bainbridge & Bownes 1981) and showed significant levels of excessively melanised tissue in the fat body, which was likely responsible for the observed developmental lethality (Figure 5C).

Conditional expansion of the *Hml*-positive cell population also increased melanisation during larval stages. RU-fed larvae showed significantly higher melanisation scores relative to the control diet-fed larvae (Figures 5D and 5E). For this purpose, we single-housed larvae to determine their genotype retrospectively (using *If* as a marker to genotype flies; see methods). As in our earlier experiment, no male larvae containing both *HmlΔ*-*GeneSwitch* and *UAS*-*ras85D^V12^* that had been fed on RU-containing media successfully developed and eclosed at 29°C. Furthermore, the genotypic proportions of pupae which eclosed were approximately the same as observed when larvae were not kept in individual vials, ruling out the possibility that the lethality phenotype was a result of competition between larvae of different genotypes (Figure 5A compared with Figure 5D). Overall, these results demonstrate that although no lifespan or infection survival phenotypes are observed upon conditional *Hml*-positive cell number modulation, manipulation of *Hml*-positive cell number using *HmlΔ*-*GeneSwitch* can produce strong functional haemocyte- specific effects.

## Discussion

Our work demonstrates that a constitutive haemocyte-specific driver (*HmlΔ*-*Gal4*) is able to not only reduce *Hml*-positive cell numbers, but also to expand the population, via expression of *bax* and *ras85D^V12^* transgenes, respectively (Figures 1A and 1B). Additionally, a conditional *HmlΔ*-*GeneSwitch* driver produced comparable effects (Figure 2). However, bleeding haemocytes from adults and larvae suggested that we were not affecting all immune cells and that compensatory mechanisms can impact the overall population *in vivo* (Supplementary Figure 2, 3). We confirmed that *Hml*- positive cell number expansion modulated a known haemocyte-dependent phenotype (self-encapsulation; Figure 5). However, manipulating *Hml*-positive cell populations had no effect on lifespan (Figure 3) or survival post-immune challenge with two different pathogens (Figure 4), in spite of a lifespan modulation being initially observed via a constitutive *HmlΔ*-*Gal4*-mediated approach (Supplementary Figure 4). No difference was observed between the strength of the constitutive and conditional *HmlΔ* drivers in terms of lethality as a result of expanding the haemocyte population (comparison between counts in Figures 1C and Supplementary Figure 7), thus reducing the likelihood that no effect was observed using our conditional *HmlΔ* as a result of it being weaker than *HmlΔ*-*Gal4*. These negative findings on longevity and especially post-infection survival are contrary to a number of studies in the literature. There are several reasons for these differences that we discuss below, with the foremost reason being that other *Hml*-negative immune cells likely replace *Hml*-positive haemocytes on ablation with *bax*.

The literature suggests that haemocyte ablation should affect survival following pathogen exposure (Shia et al. 2009; Charroux & Royet 2009; Arefin et al. 2015; Defaye et al. 2009). However, the severe pupal lethality and increased self-encapsulation phenotype observed in flies in which the *Hml*-positive population was expanded in the context of the *tuSz^1^* mutation suggests that increasing *Hml*-positive cell number really does upregulate haemocyte activity. Note, however, that we cannot distinguish cell number from other possible pleiotropic effects of expressing constitutive-active Ras, though Lemaitre and colleagues found limited evidence of activation of stress-associated pathways via overlapping approaches to manipulate haemocytes (Ramond et al. 2020); similar considerations hold for *bax* as an experimental strategy. Regardless, it is surprising that we did not observe other adult phenotypes. Although the infected flies in our experiments died within a similar timeframe as prior experiments, it is possible that the dose of the immune challenge masks any modulating effect from haemocytes. Potentially therefore, haemocyte number may be critical in the face of infection only at certain levels of infection.

Another explanation is that there is compensation within the population of cells. It is also possible that the *Hml*-negative cells are sufficient to protect against infection and that the expansion in adults via Ras expression does not reach a threshold to exert harmful or beneficial effects. It is also now known that haemocytes communicate with cross-talk between these cells and other tissues (Yang & Hultmark 2016; Green et al. 2018). As such it remains possible that both ablation and expansion of the *Hml*-positive population functionally does little, except if cells are falsely attracted to ‘self’. Perhaps, the expanded cells are differentiated such that functioning is increased in recognition of ‘self’ rather than any other functions.

Indeed, a recent study used *hid* and *reaper* to ablate *Hml*-positive haemocytes and found an increase in *Hml*-negative plasmatocytes, in line with our data and interpretation (Shin et al. 2020) and expansion of lamellocyte-like cells on ablation was also seen by Arefin and colleagues (Arefin et al. 2015). These *Hml*-negative haemocytes were shown to express *Pxn*, a classical haemocyte marker gene (Evans et al. 2014), despite the lack of *Hml*, and therefore it would be beneficial in future work to test whether our *Hml-*negative haemocyte population also expresses *Pxn*, in order to fully confirm that our results here identified the same *Hml*-negative population. Such populations of *Hml*-negative immune cells therefore offer an explanation for the lack of infection phenotype in flies where the haemocyte population has been “ablated” or “expanded”. These possibilities will prove a fruitful paradigm to study immunology in the fly; an emerging theme in this area of research are functional differences within the plasmatocyte lineage of haemocytes (Shin et al. 2020; Coates et al. 2021).

It is interesting that *ras85D^V12^* did dramatically increase the total cell population in larvae, but not in adults. This is an intriguing discovery, which suggests that potentially a feedback mechanism exists in adult flies to restrict numbers of *Hml*- positive cells which are not present or as active in larvae. This is a discovery which is not only relevant for fly biology, but also for potential use as a cancer model.

Feedback mechanisms that regulate the number of immune cells are highly relevant to understanding immunoproliferative disorders such as leukaemia or lymphoma (Connors et al. 2020; Tebbi 2021). Of further relevance, recent work from the Giangrande lab revealed early wave haemocytes can impact later blood cell development via extracellular matrix deposition, demonstrating potential feedback mechanisms are relevant in the fly (Monticelli et al. 2024).

Flies in which haemocytes are ablated display melanotic spots, purportedly as a result of the lamellocyte population expanding upon *HmlΔ*-positive cell ablation (Defaye et al. 2009; Arefin et al. 2015). We observed similar melanotic spots when we albated the *Hml*-positive cell population with the constitutive *HmlΔ*-*Gal4* driver (data not shown). Furthermore, we observed an increase in melanotic spots in the context of the *tuSz^1^* mutation when the haemocyte population was expanded. It is not clear how related these two melanotic phenotypes are. The ablation-related spots identified by previous work were found across the whole larval body (both posterior and anterior), whereas the melanisation generated by the *tuSz^1^* mutation is found only in the larval posterior (Mortimer et al. 2021). Furthermore, the *tuSz^1^*- mediated melanisation persists into adult flies, whereas the melanotic phenotype previously identified when haemocytes were ablated is only mentioned in relation to larvae (Defaye et al. 2009; Arefin et al. 2015). Although these phenotypes appear superficially similar, they may in fact originate via different mechanisms.

The flies utilised in this body of work were not grown in a sterile environment, and so our expectation was that modulation of haemocyte number would affect lifespan. Prior studies, have shown other adult phenotypes for haemocyte ablation, and perhaps if we could ablate more or all *Hml*-positive cells a phenotype would be observed, although very few *Hml*-positive cells were left in flies in which we ablated this cell population. Indeed, the *Hml*^P2A^ driver is suggested to be stronger and active in a larger fraction of the hemocyte population (Stephenson et al. 2022). However, previous uses of the *HmlΔ* promoter have shown clear immune phenotypes upon challenge with pathogens (Shia et al. 2009; Charroux & Royet 2009; Arefin et al. 2015; Defaye et al. 2009; Ayyaz et al. 2015). The only obvious difference in the approach we have used here compared to previous constitutive (or temperature-sensitive conditional) approaches is that our driver only starts to become active upon the first instance of larval feeding, as the chemical required to activate the GeneSwitch construct was in our experiments administered via the media. This should have a limited impact, considering that *Hml* is considered to only be active from early larval stages (Charroux & Royet 2009), however there is the potential for a narrow window at which *Hml* becomes active, so promoting transcription of the GeneSwitch construct, but before RU is ingested via first larval feeding. In essence this means that *HmlΔ*-positive cells may be present for a short period in our conditional haemocyte manipulation system where they would not be present when using a constitutive driver. Similarly, levels of RU-dependent activation may drop during pupal stages when active feeding has ceased. Overall, it seems unlikely that all previously documented immune phenotypes of ablating haemocytes are the result of ablation at this very early larval or pre-larval stage, and that any future ablation has no immune effect, but it cannot be ruled out.

Overall our work demonstrates that a conditional *HmlΔ*-*GeneSwitch* driver is able to drive not only ablation of *Hml*-positive cells, but also expansion of this population in flies. However, modulating haemocyte numbers conditionally did not affect longevity, in contrast to our observations when using a constitutive driver, nor did it affect immunity. Our study therefore illustrates the utility of the *HmlΔ*-*GeneSwitch* driver to understand macrophage behaviour in the fly model. The work also suggests powerful mechanisms of regulation for overall numbers of immune cells in the fly that may not always be trivial to overcome experimentally. In this process we identified potentially important but unknown mechanisms that regulate total immune cell numbers, for which conditional and especially temporal manipulation (Simons et al. 2019) using GeneSwitch provides a promising experimental paradigm.

## Acknowledgements

This work was funded by a Dunhill Medical Trust Seed Award (AIS2110\5) awarded to IE and MS, a BBSRC project grant (BB/X006603/1) awarded to IE, and a Royal Society/Wellcome Sir Henry Dale Fellowship (216405/Z/19/Z) awarded to MS. We thank Katherine Whitley and the University of Sheffield Fly Facility and Simons Lab technical team for fly husbandry and logistical support. The MDX ImageExpress Hi- content microscope is housed within the Sheffield RNAi Screening Facility and we thank Steve Brown (University of Sheffield) for assistance with that equipment. This work would not be possible without the Bloomington *Drosophila* Research Centre (NIH P40OD018537) and Flybase (MRC grant MR/N030117/1). We thank Katie Roome for contributions to initial experiments in this project and acknowledge Martin Zeidler (University of Sheffield) for helpful comments and discussions.

## Supplementary data

**Supplementary Figure 1.**
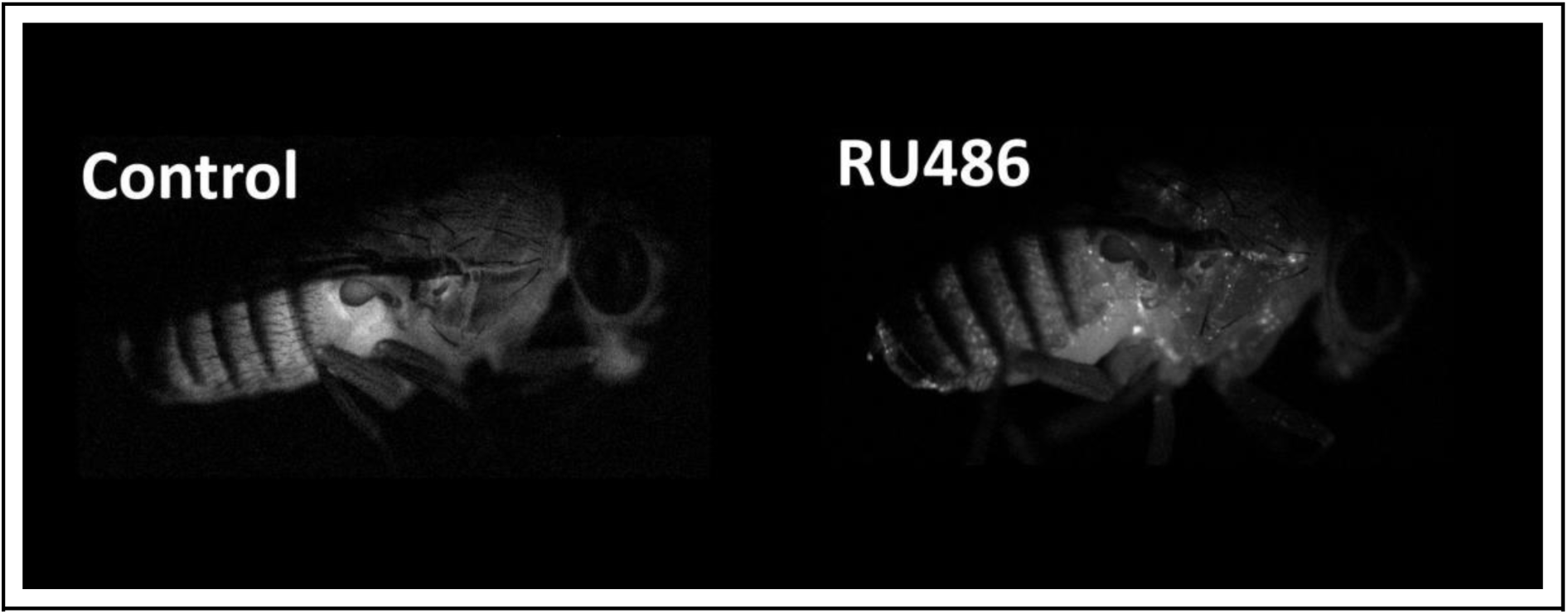
Transgenic flies carrying the *HmlΔ*-*GeneSwitch* construct were able to drive the *UAS*-*Stinger* transgene when food was supplemented with RU486, but not in its absence, and could be observed by 1 day post-eclosion. The flies used here were of the following genotype: *w^1118^*;+;*HmlΔ*-*GeneSwitch*/*UAS*-*Stinger* and were allowed to develop in the absence (‘Control’) or presence of RU486 from embryonic stages onwards (‘RU486’).

**Supplementary Figure 2.**
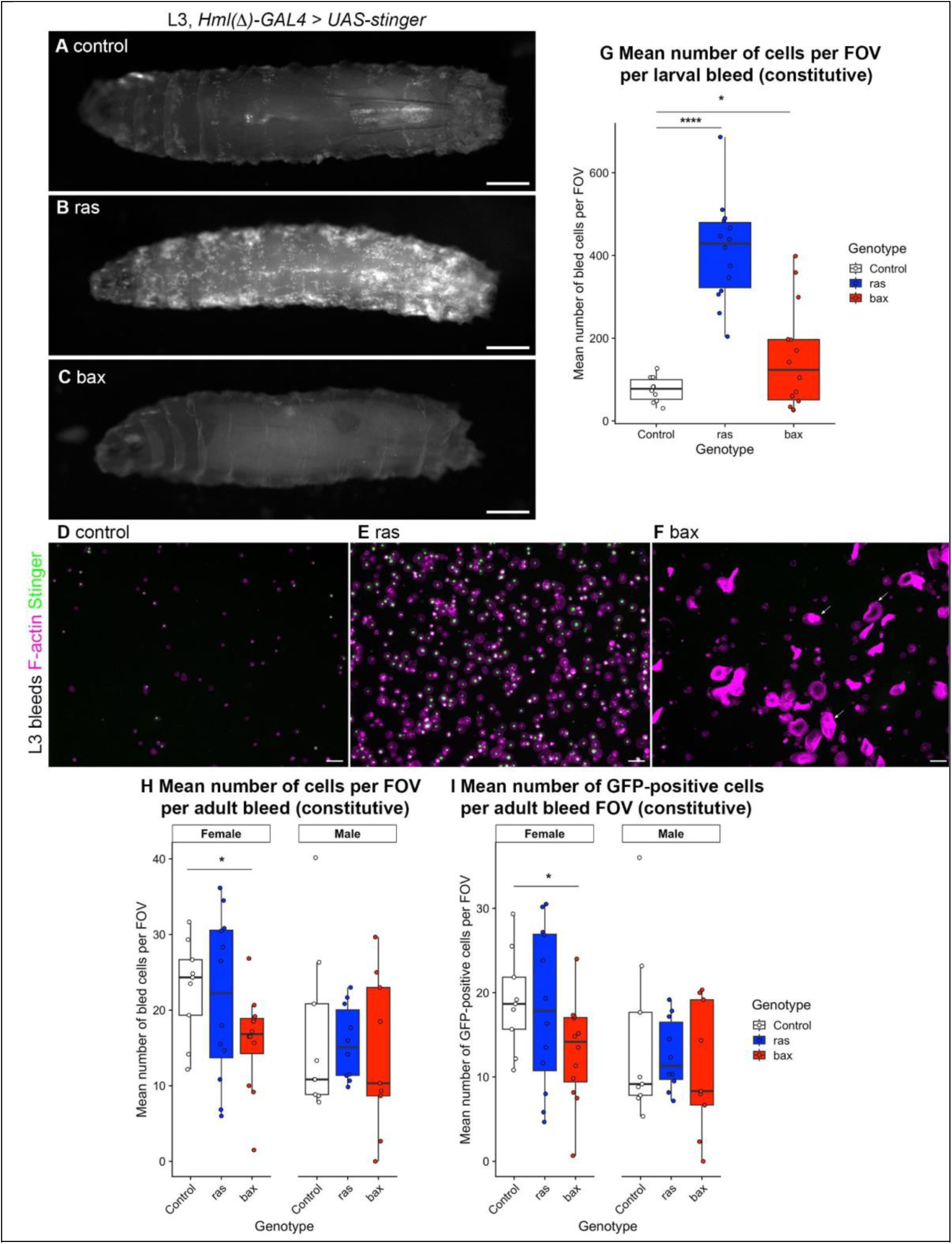
Cell counts and representative images of cells bled from larvae and adults in which Stinger and either *ras85D^V12^* or *bax* were driven by *HmlΔ- Gal4.* (A-C) L3 larvae with *HmlΔ-GAL4* used to drive constitutive expression of Stinger alone (control, A), Stinger and constitutively-active Ras85D^V12^ (ras, B), and Stinger and Bax (bax, C). GFP channel images show expression from *UAS-Stinger*; scale bars represent 500 *μ*m; anterior is left. (D-F) Representative fields of view (FOV) of haemocytes dissected from control (D), ras (E) and bax (F) L3 larvae. Merged images show F-actin (magenta) and Stinger (green) fluorescence; arrows indicate lamellocyte-like cells in bax bleeds (**F**); scale bars represent 50 *μ*m. (**G**) scatterplot showing average number of cells per FOV for L3 larval bleeds. (**H-I**) scatterplots showing average number of cells (**H**) and average number of GFP-positive cells (i.e., marked via *HmlΔ-GAL4*, **I**), per FOV, per adult dissection; n.b., two flies dissected per well. For all bleed data, each point represents the total cells bled from 1 fly or 1 larva. Statistical significance was calculated using Student’s two-tailed *t*-tests; * and **** represent p- values ≤ 0.05 and 0.0001, respectively. All genotypes are as follows: *w1118;;HmlΔ-GAL4/UAS-Stinger* (control)*, w1118;+/UAS-ras85DV12;HmlΔ-GAL4/UAS-Stinger* (ras)*, w1118;+/UAS-bax;HmlΔ-GAL4/UAS-Stinger* (bax).

**Supplementary Figure 3.**
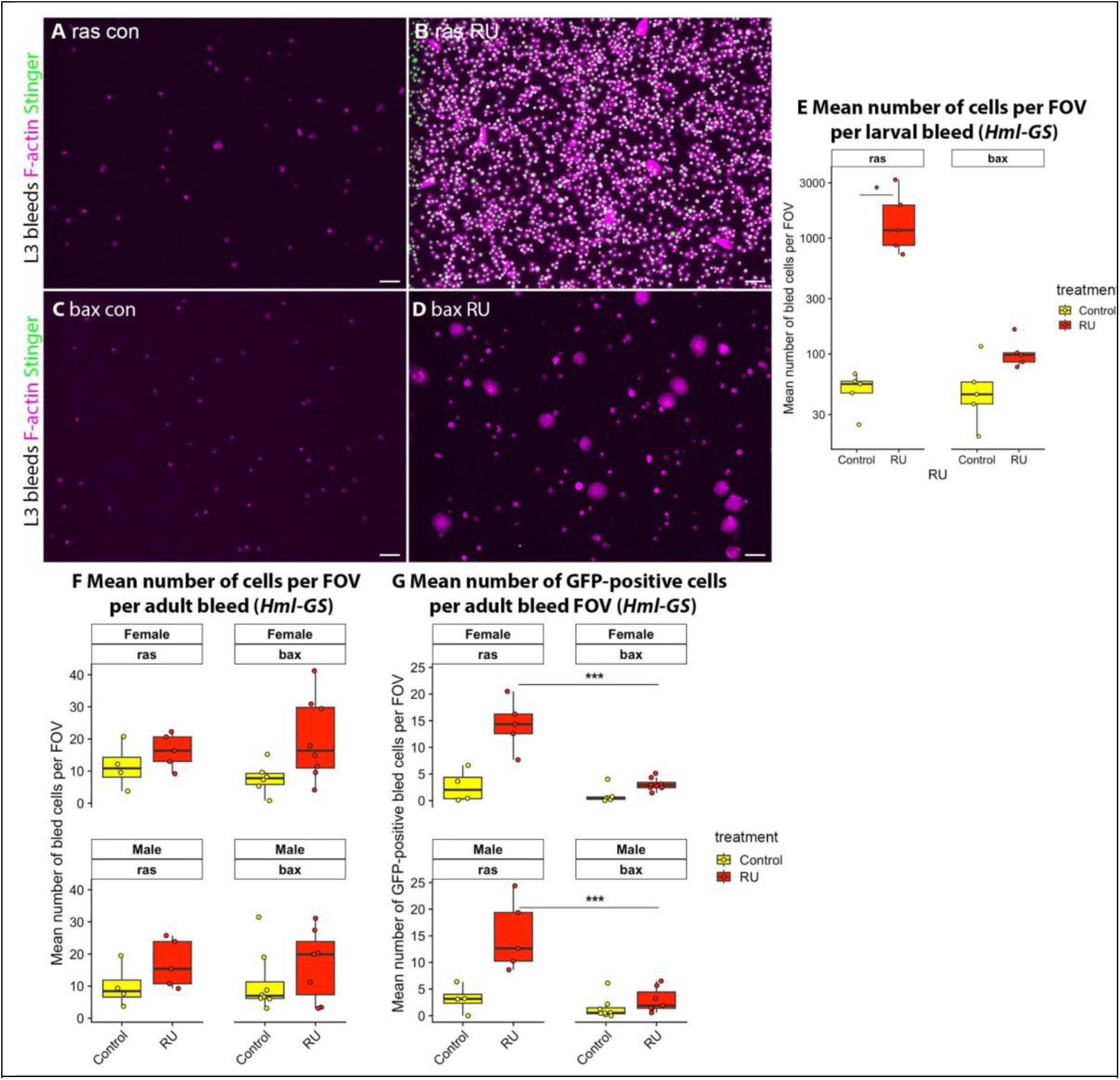
Cell counts and representative images of cells bled from larvae and adults in which Stinger and either *ras85DV12* or *bax* were driven by *HmlΔ- GeneSwitch*. (A-D) Haemocytes were bled from L3 larvae containing *HmlΔ-GS* and *UAS-Stinger,* and either *UAS-ras85DV12* (constitutively-active variant, A-B) or *UAS-bax* (C-D). Larvae were grown in the absence of RU486 as a control (ras con/bax con; A, C), or the presence of RU486 to induce expression (ras RU/bax RU; B, D). Images show F-actin (magenta) and Stinger (green) fluorescence; scale bars represent 50 *μ*m. (E) scatterplot showing average number of cells per field of view (FOV) for L3 larval bleeds with and without induction via *HmlΔ-GS* and RU486. (F-G) scatterplots showing average number of cells (F) and average number of GFP-positive cells (i.e., marked via expression from *UAS-stinger*, G), per FOV, per adult dissection; n.b., two flies dissected per well. For all bleed data, each point represents the total cells bled from 1 fly or 1 larva. Statistical significance was calculated using ANOVAs, with pairwise comparisons assessed using Tukey’s HSD post-hoc testing; * and *** represent p-values ≤ 0.05 and 0.001, respectively. All genotypes are as follows: *w1118;+/UAS- ras85DV12;HmlΔ-GAL4/UAS-Stinger* (ras con/ras RU)*, w1118;+/UAS-bax;hmlΔ-GAL4/UAS-Stinger* (bax con/bax RU).

**Supplementary Figure 4.**
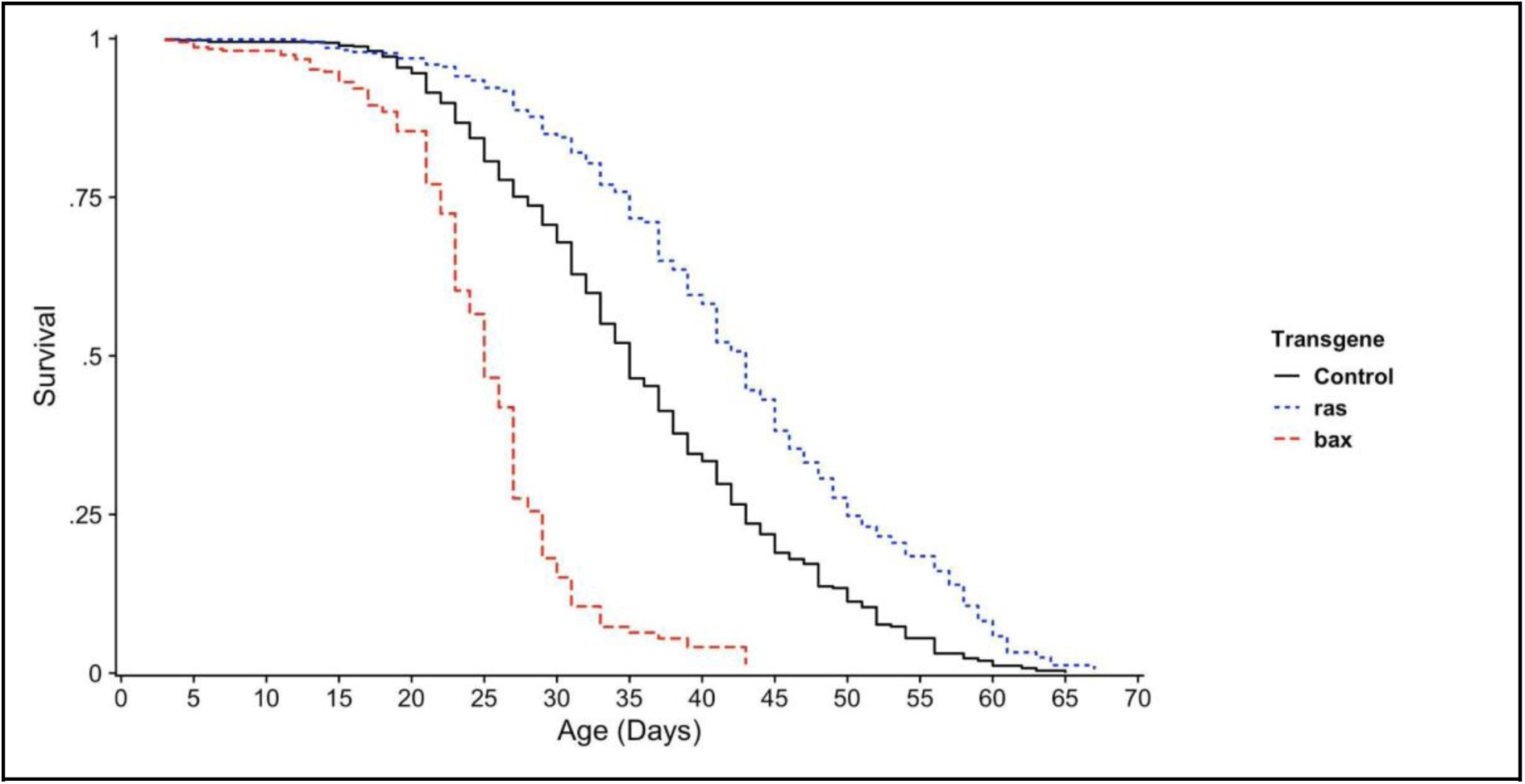
Flies carrying *bax* or *ras85DV12* transgenes show a lifespan modulation in concert with *HmlΔ*-*Gal4*. Following eclosion, flies in which *HmlΔ*-*Gal4* was used to drive *UAS*-*bax* or *UAS*-*ras85DV12* constitutively were mated for 48 hours. Subsequently flies were sorted into lifespan cages, for which new food was provided every 48 hours and the number of dead flies was counted. For ’Control’ flies, *HmlΔ*-*Gal4* drove only the *UAS*-*Stinger* transgene, whereas for ’bax’ and ’ras’ flies, either *UAS*-*bax* or *UAS*-*ras85DV12* were driven in addition to *UAS*-*Stinger*. N ≥ 360 flies for each condition. Using proportional cox hazard models, this was highly statistically significant for each transgene relative to the control (p-values < 1 x 10-20). The exact genotypes used above were as follows: ’Control’ = *w1118*;+;*UAS*- *Stinger*/*HmlΔ*-*Gal4*; ’ras’ = *w1118*;+/*UAS*-*ras85DV12*;*UAS*-*Stinger*/*HmlΔ*-*Gal4*; ’bax’ = *w1118*;+/*UAS*-*bax*;*UAS*-*Stinger*/*HmlΔ*-*Gal4*.

**Supplementary Figure 5.**
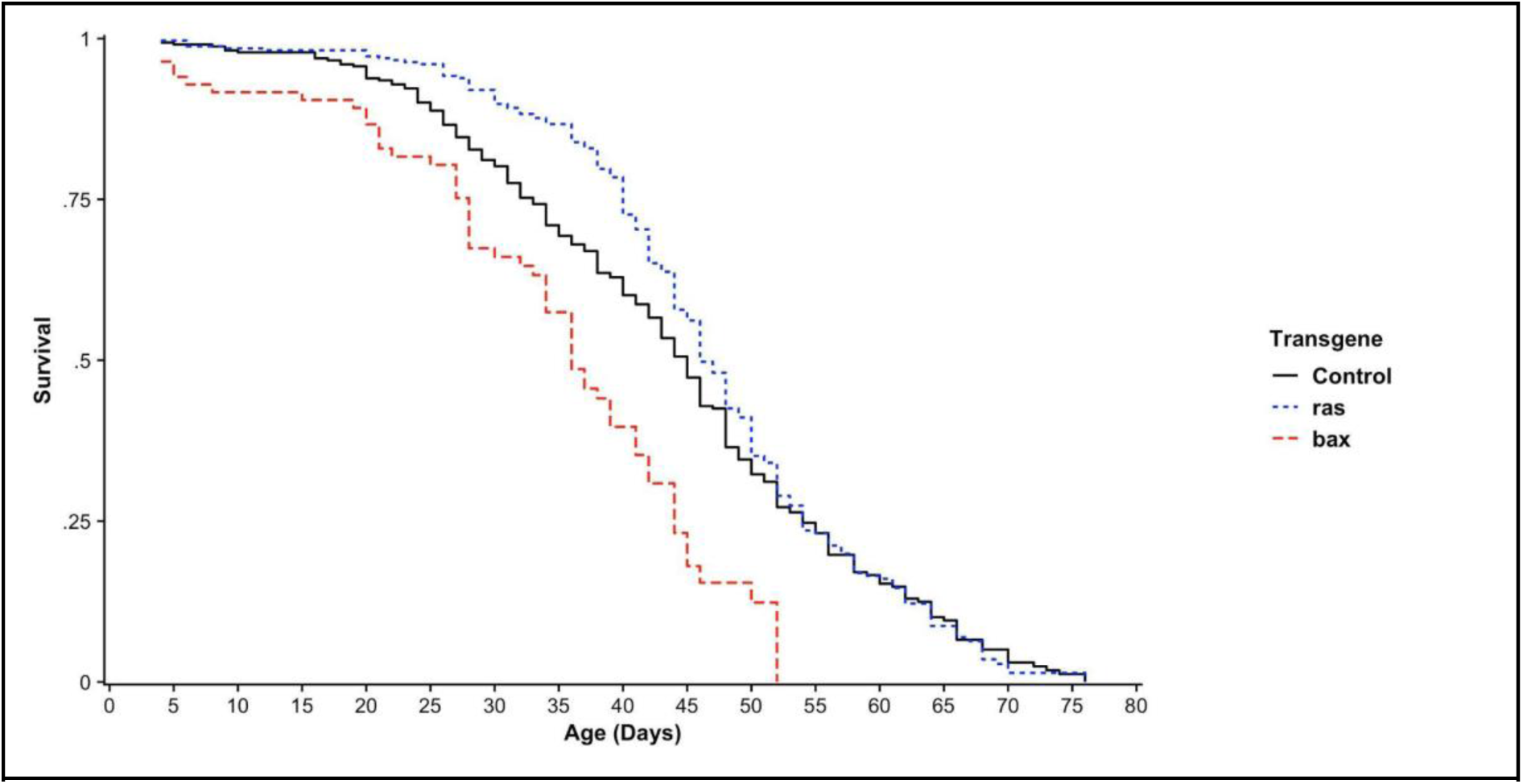
Flies carrying *bax* or *ras85DV12* transgenes show a lifespan modulation even in the absence of a *Gal4* driver. Male *UAS*-*bax*, *UAS*-*ras85DV12* or control lines were crossed to *yw* females. Progeny were mated for 48 hours and subsequently sorted into lifespan cages, for which new food was provided every 48 hours and the number of dead flies was counted. N ≥ 80 flies for each condition. P-value (bax) = 0.00002, p-value (ras) = 0.37. The genotypes used here to assess lifespan were as follows: ’Control’: *w1118*/*wyw*;+;*UAS*- *Stinger*/+; ’ras’: *w1118*/*wyw*;*UAS*-*ras85DV12*/+;*UAS*-*Stinger*/+; ’bax’: *w1118*/*wyw*;*UAS*-*bax*/+;*UAS*- *Stinger*/+.

**Supplementary Figure 6.**
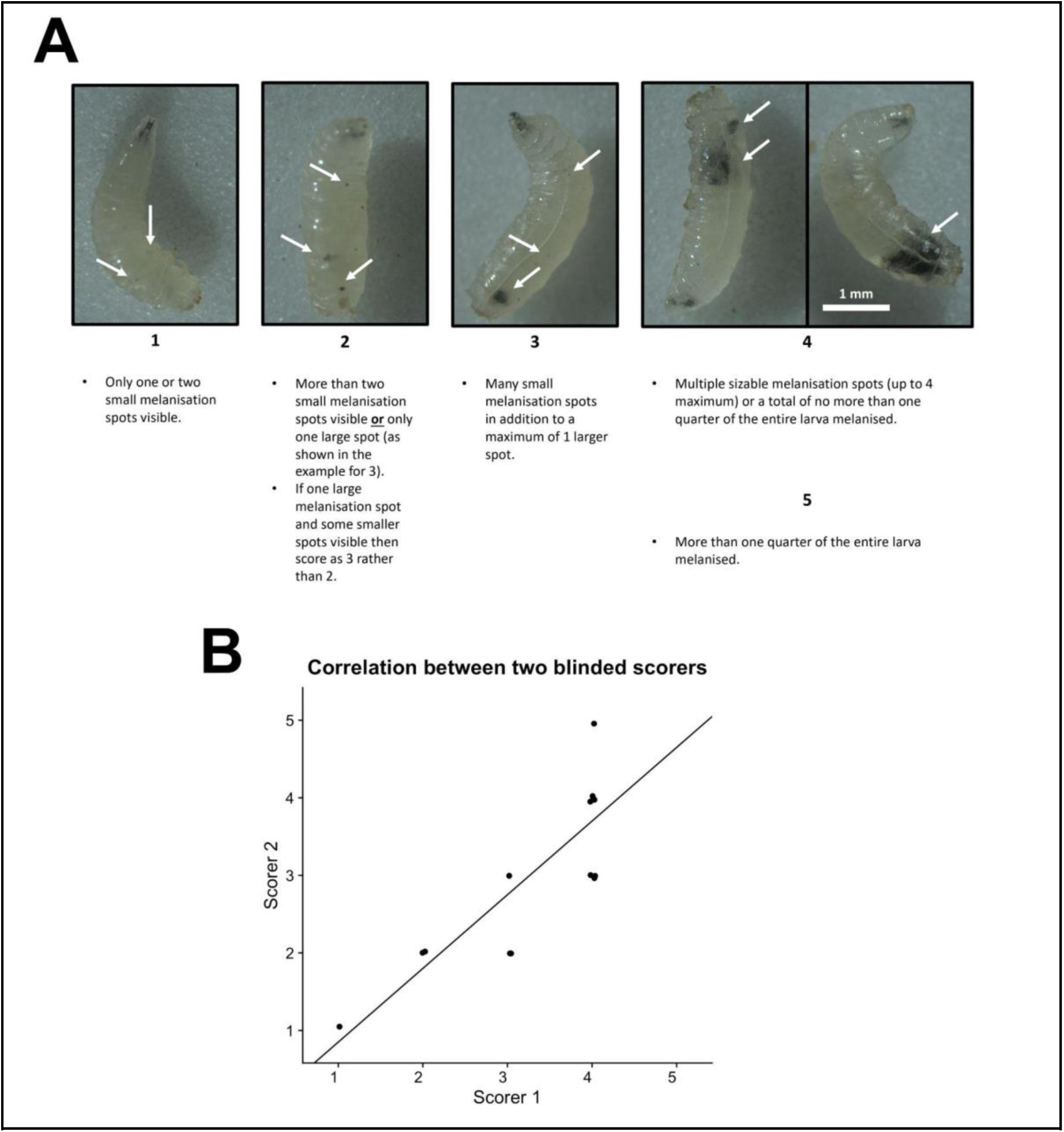
A novel semi-quantitative larval melanisation scoring system. (**A**) A novel melanisation scoring system was devised, for which L3 larvae showing signs of melanisation are picked and scored from 1-5 based on the displayed criteria. Although every male larva displayed some level of melanisation in the experiments undertaken here, the scoring system could conceivably be extended to include a 0 score as well, to signify absolutely no excessive melanisation at all if required. Examples of each score are also shown, with white arrows indicating notable features of each score. 2 examples of larvae, both of which would be scored as 4 are shown. In the experiments undertaken here, no larvae were severe enough to be given a score of 5, but the criterion for a score of 5 is also shown. (**B**) A blinded correlation of scores given to 15 larvae by 2 scorers using the melanisation scoring system. A Spearman correlation test resulted in ρ = 0.856 and p-value = 4.72 x 10-5, signifying a good correlation between the two independent sets of scores given. A small random value (- 0.05 to 0.05) is added to each score on the correlation plot, for the purposes of illustrating the individual points present, but these values were not used for the statistical analysis. All larvae used to produce this larval melanisation scoring system contained the *tuSz1* X chromosome mutation.

**Supplementary Figure 7.**
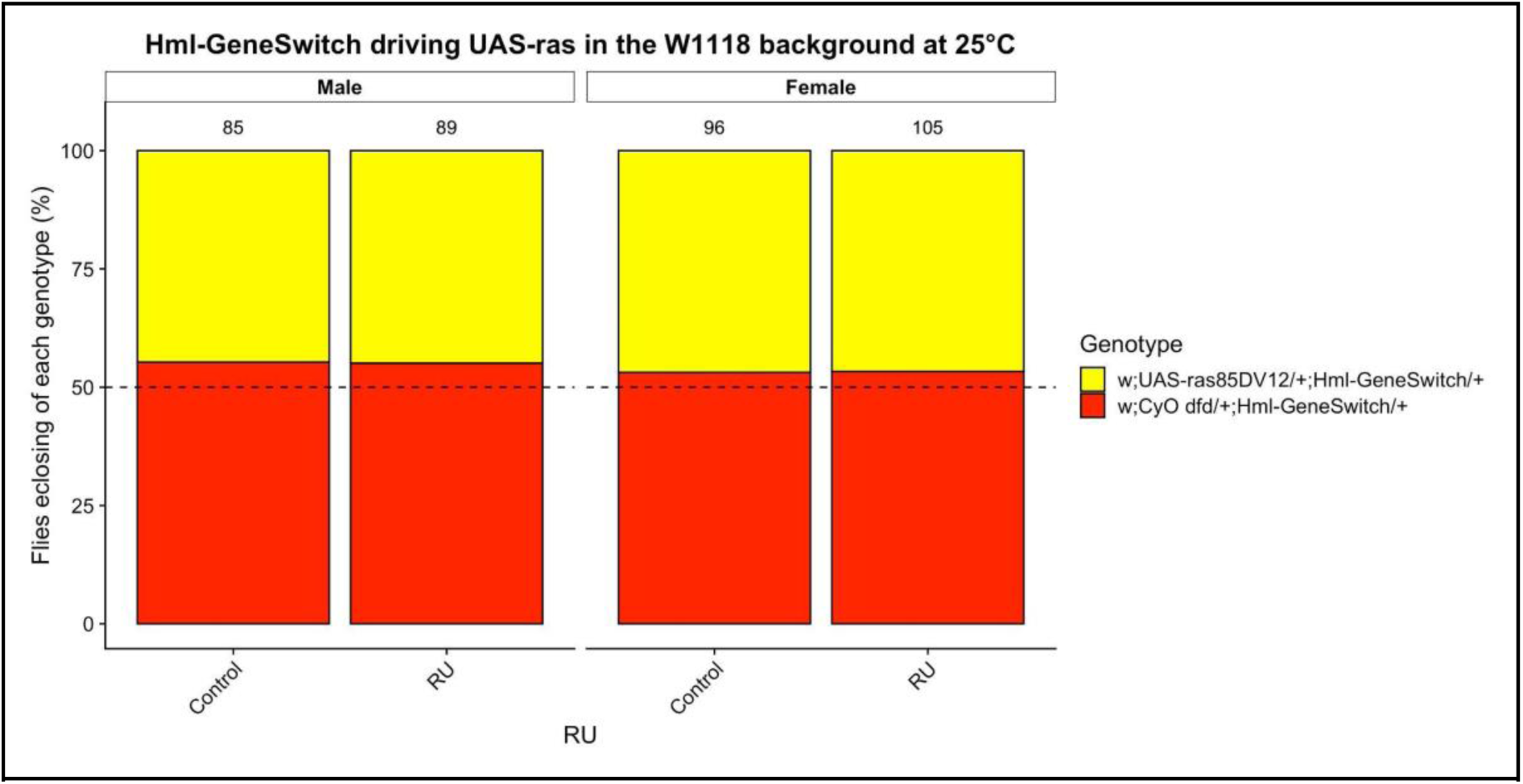
Eclosion ratios in a control genetic background. Female *w1118;+;HmlΔ-GeneSwitch* flies were crossed with male *w1118;UAS-ras85DV12/CyO dfd* at 25°C. The number of flies eclosing with *CyO dfd* was compared to those eclosing without it in either control or RU conditions, so as to genotype the eclosing flies. The estimated ratio for each condition was 50%, indicated by the dotted line. The total number of progeny counted from each cross is shown above each bar.

**Supplementary Figure 8.**
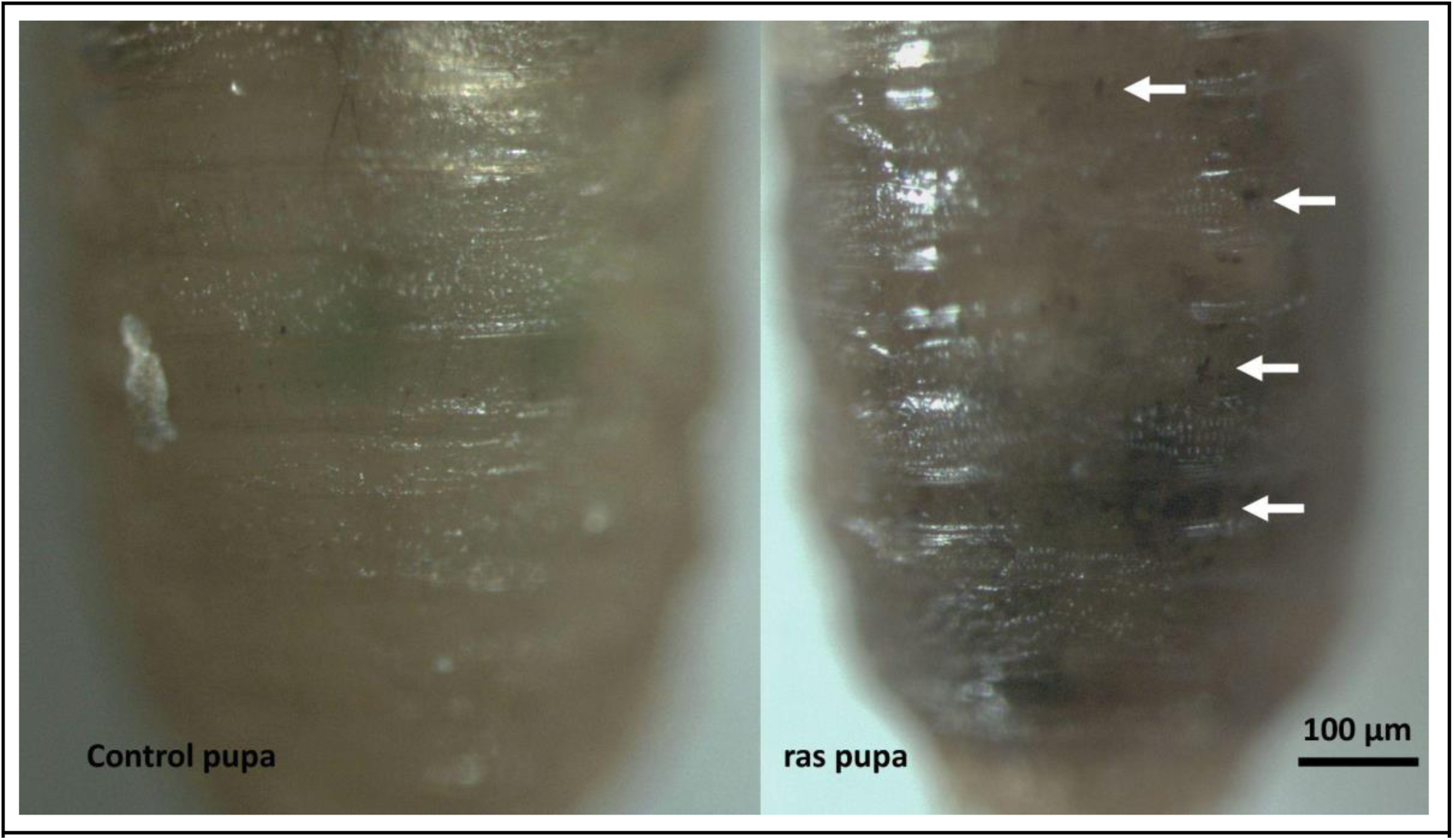
Pupae in which the *Hml*-positive cell population had been expanded using *HmlΔ-GeneSwitch* (“ras pupa”) showed high levels of excessive melanisation compared to control pupa, which was clear even through the puparium. White arrows indicate clear melanotic spots in the example ras pupa.

**Supplementary Table 1.** Statistics for the effect of *Hml*-positive cell ablation and expansion on lifespan using *HmlΔ*-*GeneSwitch*. HR is the hazard ratio. All statistics calculated using the Coxme R package. Each sample is a single fly.

| Condition | Sex | RU vs control statistics |  |  | Sample sizes |  |
| --- | --- | --- | --- | --- | --- | --- |
|  |  | ln(HR) | se(HR) | P-value | Control | RU |
| Control | Female | 0.865 | 0.081 | 0.075 | 827 | 554 |
| bax | Female | 0.949 | 0.094 | 0.577 | 282 | 253 |
| ras | Female | 1.074 | 0.081 | 0.381 | 404 | 335 |
| Control | Male | 1.110 | 0.060 | 0.083 | 685 | 729 |
| bax | Male | 0.943 | 0.088 | 0.509 | 270 | 313 |
| ras | Male | 0.865 | 0.081 | 0.074 | 401 | 335 |

**Supplementary Table 2.**
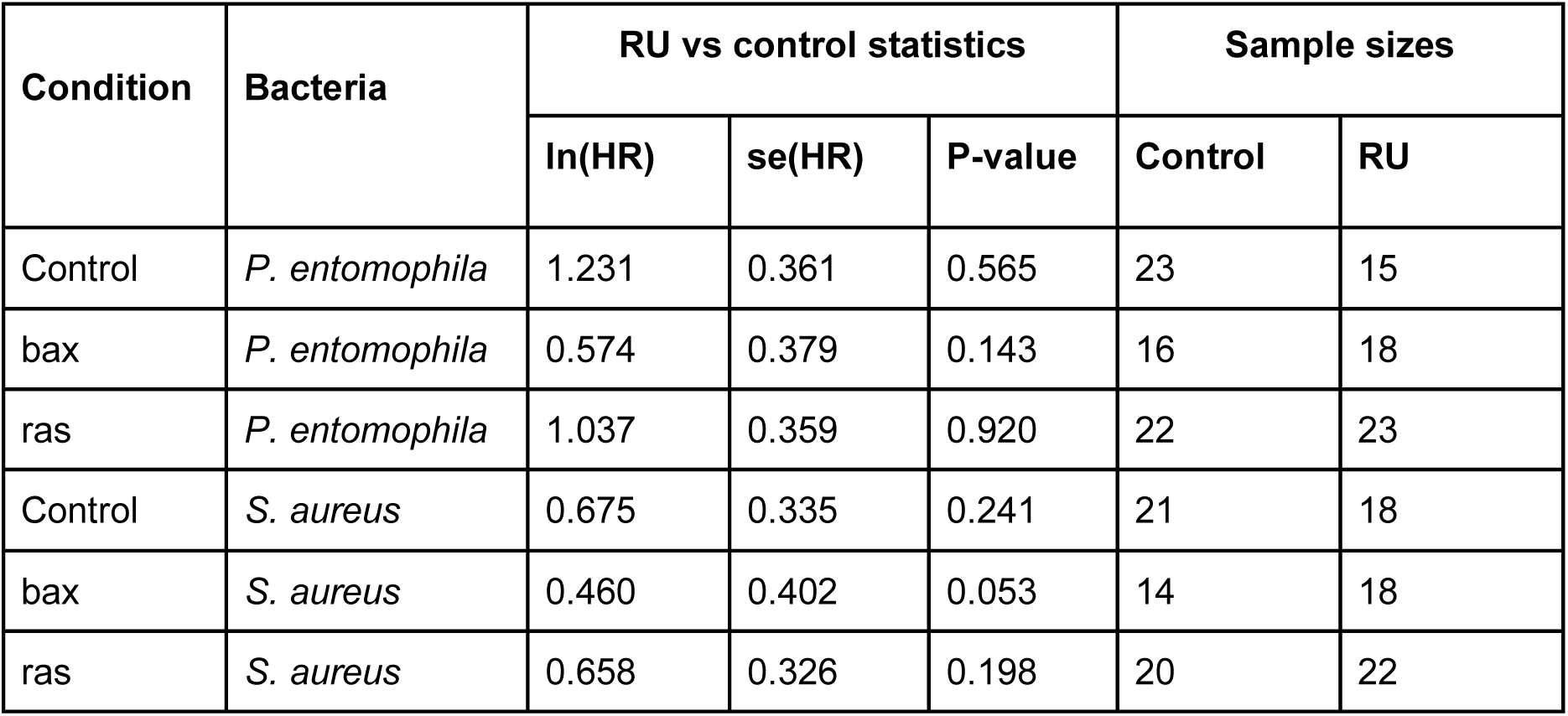
Statistics for the effect of *Hml*-positive cell ablation and expansion on post-infection survival using *HmlΔ*-*GeneSwitch*. HR is the hazard ratio. All statistics calculated using the Coxph R package. Each sample is a single fly.

